# Moving bar of light evokes vectorial spatial selectivity in the immobile rat hippocampus

**DOI:** 10.1101/2021.12.28.474337

**Authors:** Chinmay S. Purandare, Shonali Dhingra, Rodrigo Rios, Cliff Vuong, Thuc To, Ayaka Hachisuka, Krishna Choudhary, Mayank R. Mehta

**Affiliations:** Department of Physics and Astronomy, W.M. Keck Center for Neurophysics, University of California at Los Angeles, Los Angeles, CA 90095, USA; Department of Bioengineering, University of California at Los Angeles, Los Angeles, CA 90095, USA; Department of Neurology, Department of Electrical and Computer Engineering, University of California at Los Angeles, Los Angeles, CA 90095, USA

## Abstract

Visual cortical neurons encode the position and motion direction of specific stimuli retrospectively, without any locomotion or task demand^1^. Hippocampus, a part of visual system, is hypothesized to require self-motion or cognitive task to generate allocentric spatial selectivity that is scalar, abstract^2,3^, and prospective^4–7^. To bridge these seeming disparities, we measured rodent hippocampal selectivity to a moving bar of light in a body-fixed rat. About 70% of dorsal CA1 neurons showed stable activity modulation as a function of the bar’s angular position, independent of behavior and rewards. A third of tuned cells also encoded the direction of revolution. In other experiments, neurons encoded the distance of bar, with preference for approaching motion. Collectively, these demonstrate visually evoked vectorial selectivity (VEVS). Unlike place cells, VEVS was retrospective. Changes in the visual stimulus or its trajectory did not cause remapping but only caused gradual changes. Most VEVS tuned neurons behaved like place cells during spatial exploration and the two selectivity were correlated. Thus, VEVS could form the basic building block of hippocampal activity. When combined with self-motion, reward, or multisensory stimuli^8^, it can generate the complexity of prospective representations including allocentric space^9^, time^10,11^, and episodes^12^.

## Introduction

Sensory cortical neurons generate selective responses to specific stimuli, in an egocentric (e.g. retinotopic) coordinate frame, without any locomotion, memory or rewards^1^. In contrast, the hippocampus is thought to contain a visually evoked, abstract, allocentric cognitive map, supported by spatially selective place cells^2^, grid cells^13^ and head direction cells^14^. These responses are thought to arise from not only visual^3^ but self-motion cues^9,15^ too, e.g. via path integration^16^. Recent studies demonstrate hippocampal activity modulation by auditory^17–19^ or social stimuli^20,21^. However, to elicit these responses, additionally behavioral, cognitive or reward variables were required, whose removal nearly eliminated hippocampus selectivity^17,18,22–25^. No study has demonstrated hippocampal neural selectivity to a moving visual stimulus, without self-motion or rewards, similar to that found in visual cortices, even though they provide major input to the hippocampus, and are thought to be crucial for hippocampal function. Hence, we investigated if place cells encode the angular and linear position as well as motion direction of a simple stimulus, regardless of self-motion, memory or reward.

Rats were gently held in place on a large spherical treadmill, surrounded by a cylindrical screen^26^. They could move their heads around the body by a small amount but couldn’t turn their body. To keep them motivated, they were given random rewards, similar to typical place cell experiments, and were pretrained to do a virtual navigation task in the same apparatus (See methods). The only salient visual stimulus during the experiments was a vertical bar of light, 74cm tall, 7.5cm wide, 33cm away from the rat, thus subtending a 13° solid angle. In the first set of experiments, the bar revolved around the rat at a constant speed (36°/s), without any change in shape or size (Fig. 1a, b). The bar’s revolution direction switched between CW (clockwise) and CCW (counterclockwise) every four revolutions. In subsequent experiments, we varied the color, pattern, movement direction and trajectory of the bar. A majority of neurons showed selective responses in all cases.

**Fig. 1.**
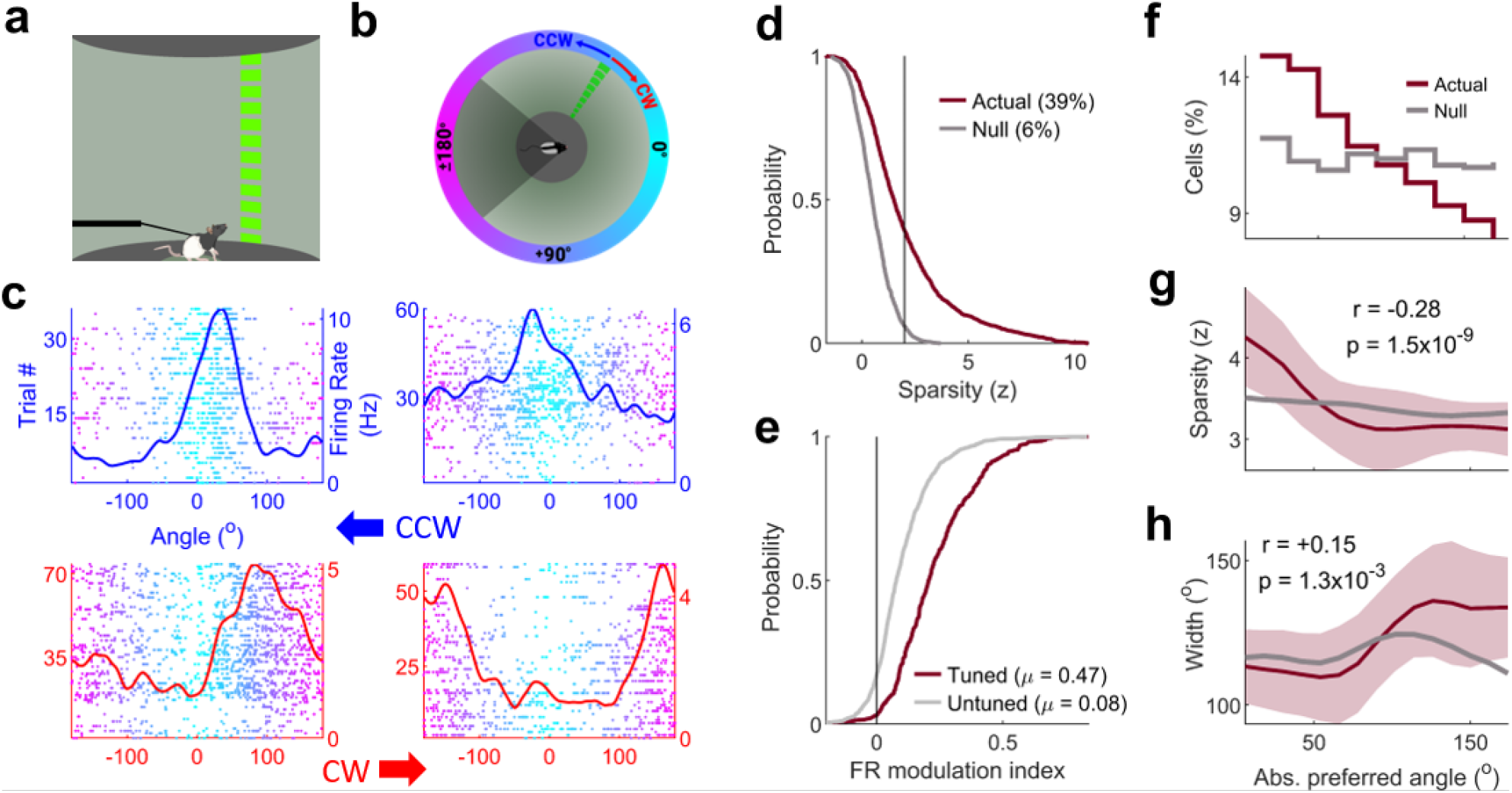
Hippocampal response to a revolving bar of light: (**a**) Experimental setup schematic and (**b**) its top-down view. Rat’s field of view is 270°. The region behind him (dark gray) is invisible to him. (**c**) Trial number (y-axis on the left) and firing rates (y-axis on the right) of 4 CA1 neurons as a function of the angular position of the bar (front= 0°, behind= ±180°). Revolution direction of the bar: Top panels, blue, counterclockwise (CCW); bottom panels, red, clockwise (CW)). (**d**) Cumulative distribution function (CDF) of strength of tuning (z-scored sparsity, see methods); response with higher tuning chosen between CCW and CW, (d) through (f)). The measured data shows significantly greater (*p*=1.26×10^−89^, KS-test here and subsequently) tuning than the shuffled data (Gray line). 39% of neurons showed significant (z>2) tuning. (**e**) CDF of firing rate modulation index within versus outside the preferred zone (see methods) for tuned cells (*z*>2) was significantly different (*p*=1.9×10^−50^) than untuned (*z*<2) cells. (**f**) Twice as many tuned cells (y-axis) had their preferred angles (angle of maximal firing, x-axis) in the front than behind. (**g**) z-scored sparsity (solid line=Median, shaded area=SEM here and subsequently) of tuned cells decreased as a function of their preferred angle. (*Pearson* correlation coefficient, here and subsequently, in inset). (**h**) Full width at quarter maxima of tuned responses increased with preferred angle.

### Most CA1 neurons encode stimulus angle

We measured the activity of 1191 putative pyramidal neurons (with firing rate above 0.2Hz during the experiment) from the dorsal CA1 of 8 Long-Evans rats in 149 sessions using tetrodes (see methods^8^). Many neurons showed clear modulation of firing rate as a function of the angular position, i.e., aVEVS (Fig. 1c), with elevated firing rates in a limited region of visual angles. Across the ensemble of neurons, 464 (39%) showed significant (sparsity (*z*)>2, corresponding to *p* < 0.023, see methods, Extended Data Fig. 1) tuning in either the CW or CCW direction (Fig. 1d). These were classified as tuned cells, in contrast with untuned cells with z<2.

Like the visual cortical neurons and hippocampal place cells, most tuning curves were unimodal (Extended Data Fig. 2) with a single preferred angle where the firing rate was the highest. Off responses (a significant decrease in firing rate) were virtually nonexistent. The preferred angles spanned the entire range, including angles behind the rat that he could not see barring rare occasions (Fig. 1e). These responses resembled striate cortical neurons in many ways^1,27^. More neurons encoded the positions in front of the rat (0°) and there was a gradual, two-fold decline in the number of tuned cells for angles behind (±180°). The strength of aVEVS (Fig. 1f, see methods) was much larger near 0° than near 180°. The tuning curve width increased gradually (from 114° to 144° Fig. 1g) as a function of the absolute preferred angle from 0° to 180°. But, the widths were quite variable at every angle, spanning about a third of the visual field, similar to place cells on linear tracks^28,29^.

Hippocampal place cells on 1D tracks have high firing rates within the field with little spiking outside^28^. In contrast, most neurons with significant aVEVS spiked considerably outside the preferred zone, as evidenced by modest values of the firing rate modulation index (Fig. 1h, see methods). These broad aVEVS tuning curves resembled the angular tuning of CA1 neurons recently reported in the real world and virtual reality^30^, with comparable fraction of neurons showing significant angular tuning. The trial to trial variability of aVEVS was comparable to recent experiments in visual cortex of mice under similar conditions^31^. Notably, the variability in the mean firing rate across trials was small and unrelated to the degree of aVEVS. However, the trial-trial variability of the preferred angle was quite large and predictive of the degree of aVEVS of a neuron (Extended Data Fig. 3).

### Revolution dire ction selectivity of aVEVS

In the primary visual cortex, majority of neurons respond selectively to the angular position of a stimulus, regardless of its movement direction^1^. But, majority of hippocampal neurons on linear tracks are directional, with far greater firing rate in one direction of journey^8,28^. Further, in both areas, neurons that are active in both directions, show significant and stable selectivity in both directions too. Hence, we inspected the selectivity, directionality, and stability (see methods) of the aVEVS.

The firing rates were comparable for most neurons in two directions of revolution (see below). The degree of tuning varied continuously across neurons with no clear boundary between tuned and untuned neurons (Extended Data Fig. 4). Some neurons were bidirectional, i.e., significant (z>2) aVEVS in both directions of revolution (Fig. 2a, Extended Data Fig. 4). However, a larger subset of neurons was unidirectional, with significant (z>2) aVEVS in only one movement direction (Fig. 2b, Extended Data Fig. 4). Surprisingly, there were many untuned-stable neurons, which were deemed untuned based on the standard, z-scored sparsity criteria (z<2) but showed consistent, significantly stable (stability KS-test *p<*0.05, see methods) spiking across trials (Fig. 2c, Extended Data Fig. 6). Across the ensemble, 13% (154) of neurons were bidirectional, 26% (310) were unidirectional, and the majority, 35% (421) were untuned-stable (Fig. 2d, Extended Data Fig. 4). Thus, the vast majority (74%, 885) of hippocampal pyramidal neurons were consistently modulated by the angular position and direction of the revolving bar. However, unlike visual cortex, far more aVEVS neurons were unidirectional, and unlike hippocampal place cells and visual cortex, far greater number of neurons showed untuned but stable responses. The preferred tuning angle was around 0°, i.e. in front of the rat for most neurons (Extended Data Fig. 5) and this bias was greater for the bidirectional cells.

**Fig. 2.**
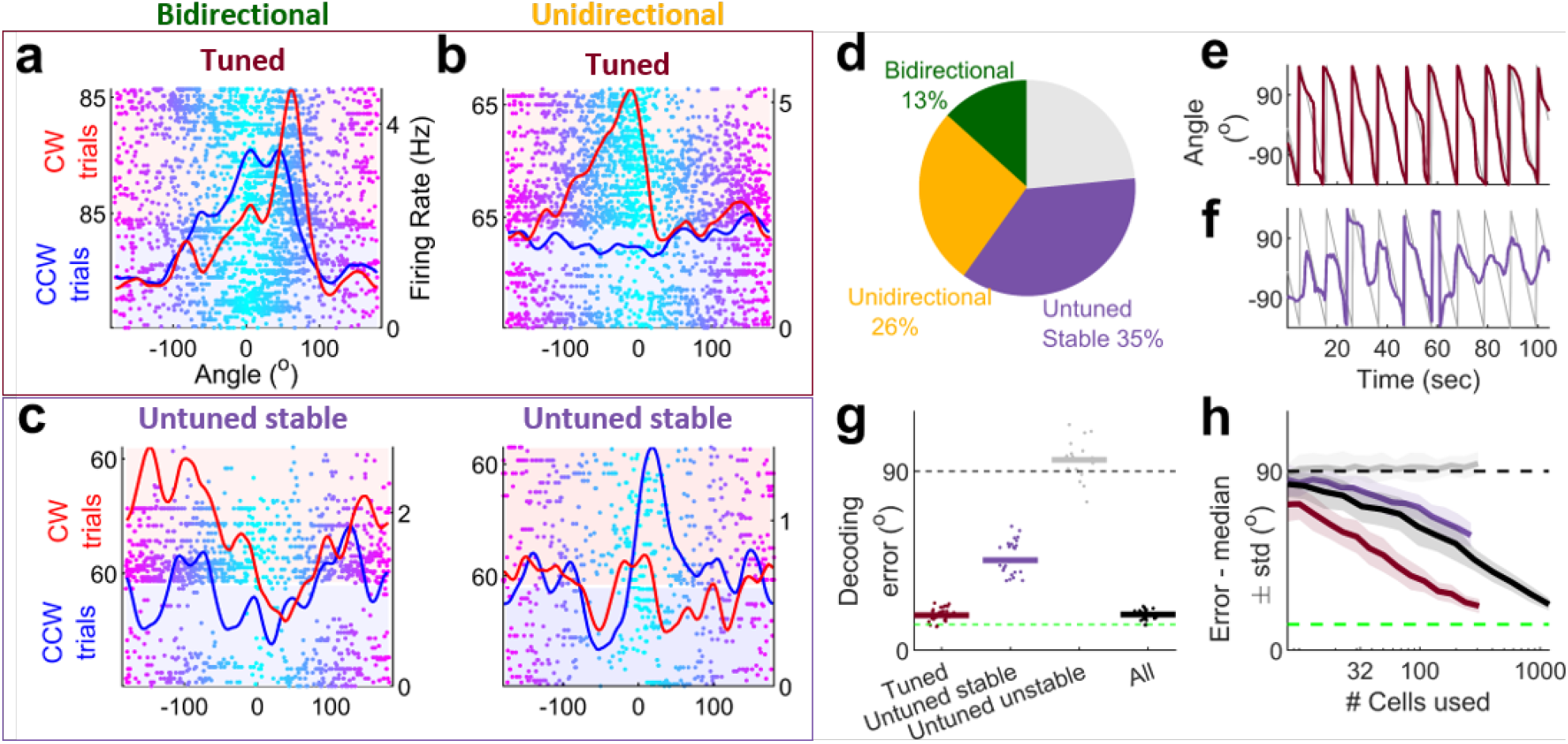
Directionality, stability and ensemble decoding of aVEVS. (**a**) A bi-directional cell, with significant (z>2) tuning (maroon) in both CCW and CW directions. (**b**) Similar to (a), but for a uni-directional cell, with significant tuning in only one direction (CW). (**c**) Cells with multi-peaked, stable responses (lavender), that did not have significant sparsity or tuning (z<2) (bi-directional stable, left; unidirectional stable (CCW), right). (**d**) Distribution of selectivity. (**e**) Decoded angle using only tuned cells in the CCW direction (maroon) **(f)** Same as (e) but using only the untuned-stable cells (lavender). (**g**) Median error between stimulus angle and decoded angle over 30 instantiations of 10 trials each for actual and shuffle data. The decoding errors for tuned (µ±sem=17.6°±0.6°) and untuned stable (45.2°±1.4°) were significantly less than that of shuffles (KS-test *p*=1.8×10^−14^ for both). Green dashed line indicates width of the visual cue; black dashed line indicates median error expected by chance. (**h**) Decoding error decreases with increasing population size for all (black), tuned (maroon) and untuned stable (lavender) cells, but not for untuned unstable cells (gray).

Highest (and significant) correlation between CCW and CW tuning curves was seen for bidirectional cells, followed by unidirectional and then untuned stable cells, but not the untuned unstable cells (Extended Data Fig. 5). The mean rates were comparable in two directions, but significantly larger in the direction with better aVEVS, largely because of an increase in firing within the preferred zone (±90° around the preferred angle) in the tuned direction. Higher rate cells were more likely to be bidirectional, than unidirectional, even after accounting for the differences in firing rate (Extended Data Fig. 6).

### Population vector de coding of aVEVS

In addition to individual cells, the population responses were also coherent for tuned and untuned-stable populations (Extended Data Fig. 7, see methods). During spatial exploration, the ensemble of a few hundred place cells is sufficient to decode the rat’s position using population vector decoding^32^. Using similar methods, we decoded the angular position of the bar (see methods).

The ensemble of 310 tuned cells (CCW), with a short temporal window of 250ms, could decode the angular position of the bar with a median accuracy of 17.6° (Fig. 2h, j) comparable to the bar width (13°) and similar to position decoding with place cells^7,32^. Additionally, the 266 untuned stable cells could also decode the position of the bar significantly better than chance. Though the median error was 45.2° (Fig. 2i, j) was larger than that for the tuned cells, it was much smaller than chance (90°), further demonstrating significant aVEVS in untuned stable cells. Decoding performance improved when using a larger number of tuned or untuned stable cells (Fig 2k). Thus the ensemble of untuned stable cells contained significant stimulus angle information, even though these individual cells did not^33^. This was not the case for the untuned unstable cells.

### aVEVS is re trospective

Under most conditions, visual cortical neurons respond to the stimulus with a short latency, i.e. retrospectively, whereas most hippocampal bidirectional cells on linear tracks are prospective^3^, i.e. they fire before the rat reaches a given position from the opposite movement directions^6–8^. However, the converse was true for the bidirectional aVEVS (Extended Data Fig. 8). The preferred angle in the CCW direction lagged behind that in the CW direction (example cell, Fig. 3c), i.e., in both directions the neuron responded to the bar after it had gone past a specific angle, which is a retrospective response. The circular difference between the preferred angle between the CW and CCW directions (bidirectional population response, Fig. 3a-b), was predominantly positive. Are only the peaks of aVEVS retrospective or do the entire tuning curves show lagged responses? The cross correlation between the entire tuning curves between the CW and CCW directions (Fig. 3d) of the majority (80%) of neurons showed maximum correlation at positive latency. Thus, most neurons responded to the oriented bar retrospectively, i.e., with a lag.

**Fig. 3.**
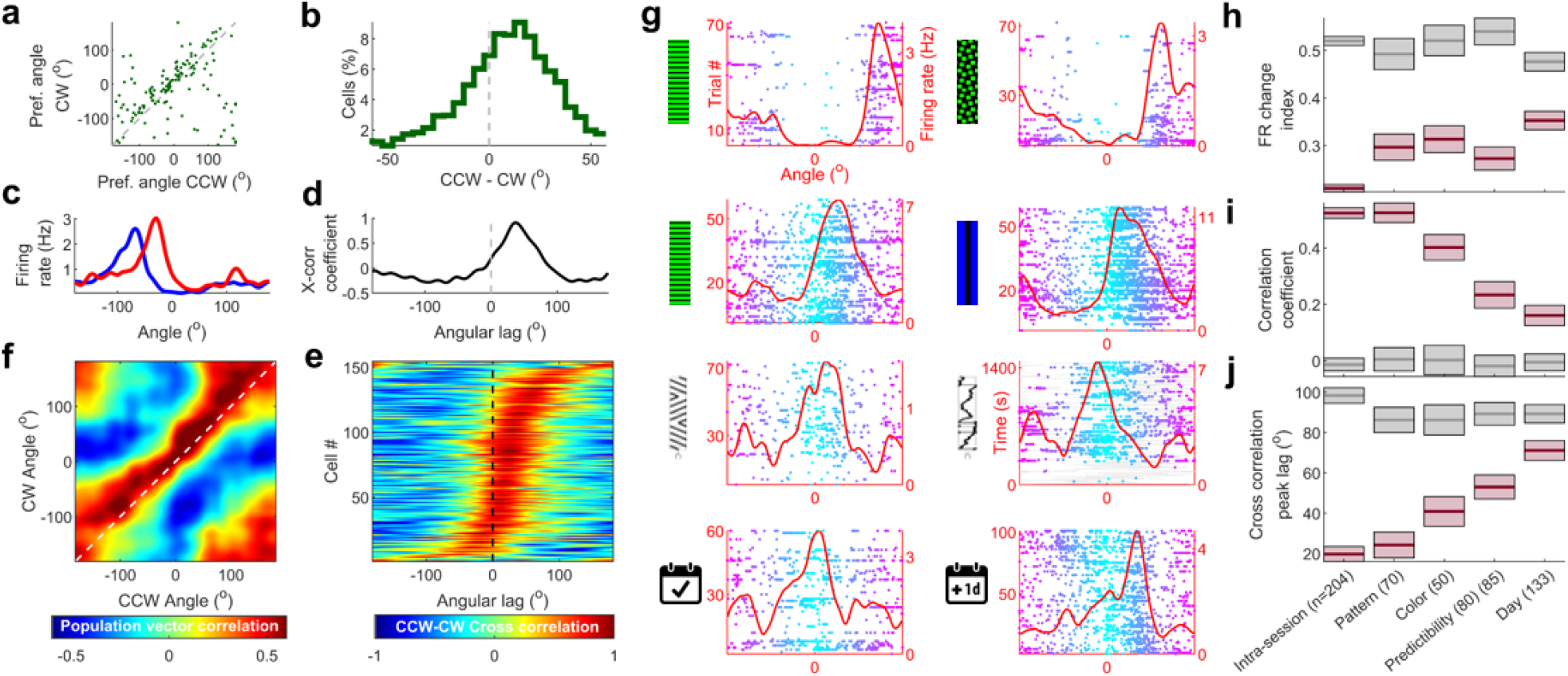
aVEVS is retrospective and changes gradually with stimulus pattern, color, motion predictability and time. (i) For bidirectional tuned cells, the preferred angle in the CW (y-axis) direction was greater than that in the CCW (x-axis). Histogram of difference (CW-CCW, restricted to ±50°) was significantly (t-test at 0°, *p*=0.003) positive indicating a retrospective shift. (**c**) Retrospective latency between the CCW (blue) and CW (red) tuning curves for a birectional cell (**d**) Cross correlation between the CCW and CW responses in (d) was maximum at positive latency (+27°). (**e**) Such cross correlations were performed for each bidirectional cell and sorted according to their peak-lag. Majority (80%) of lags were positive. (median=+19.9°±49.8°, circular median t-test at 0°, *p*=4.8×10^−16^). **(f)** Population vector overlap of aVEVS had a significant (circular median t-test, *p=*1.5×10^−36^) peak at a positive lag (median=+54.3^°^ ±25.3°). (**g**) Change in pattern (green-striped (left) vs green-checkered (right)) caused the smallest change in aVEVS. Change in color (green vs blue), and pattern (horizontal vs vertical stripes) caused gradually greater change in aVEVS. Changes in predictability of the stimulus motion, or mere passage of time (1 day) caused the greatest changes. Only CW example cells shown here. (**h)** Firing rate remapping, quantified by FR change index was significantly (KS-test, *p-*value range - 1.2×10^−90^ to 1.4×10^−5^) smaller for the actual data (dark-pink) than for shuffle data (gray) for all conditions. Thick line – median, box – sem, here and through (j). **(i)** Similar to (e), correlation coefficient between the tuning curves across different conditions was significantly greater than shuffle (KS-test *p-*value range-2.6×10^−58^ to 3.8×10^−6^). **(j)** Same as (e), angular lag in cross correlation to quantify amount of shift between tuning curves across the two conditions. All were significantly lesser than shuffle (KS-test *p-*value range-4×10^−46^ to 1.3×10^−3^), *n=*number of responses measured.

The median latency to response was 276.2ms, which translates to 19.9° median shift in cross correlation (Fig. 3e). This retrospective coding was evident across the entire ensemble of bidirectional cells, such that the population vector overlap between the two directions was highest at values slightly above the diagonal (Fig. 3f, see methods).

Unidirectional cells too showed retrospective tuning, with cross-correlations latency (19.9°, or 276.2ms) comparable to bidirectional cells (Extended Data Fig. 9). Thus, the retrospective coding does not arise due to difference in tuning strengths. Small but significant retrospective bias was also observed in the untuned-stable cells but not for the unstable cells (Extended Data Fig. 9). Additional experiments using a photodiode showed that this lag could not be explained by the latencies in the recording equipment (Extended Data Fig. 10).

### Gradual change in aVEVS with stimuli and time

Change in distal visual cues causes remapping in place cells, i.e. large changes in firing rate, degree of spatial selectivity and the preferred location^34,35^. On the other hand, primate hippocampal neurons show selectivity to a visual stimulus^36^. To address this, we measured the responses of the same set of neurons, on the same day, to bars of light with gradual changes in their visual features (see methods), without any other changes. First, we changed the stimulus minimally (pattern change, Fig. 3g row #1, h-j). Neural firing rates, preferred tuning location and tuning curve profiles were largely invariant and comparable to intra-session variation (Fig. 3h-j, see methods). Next, we introduced larger change in the bar appearance by changing both color and pattern. This resulted in significantly more changes in all measures of aVEVS, though this too was far less than expected by chance.

Sequential tasks can influence neural selectivity in the hippocampus^9,10^ and visual cortex^37^. Hippocampal neurons also show selectivity in sequential, non-spatial tasks^19–21^. Sequential versus random goal-directed paths induce place field remapping in the real world^38^ and large change in selectivity in VR. To compute the contribution of the sequential movement of the bar of light to aVEVS, we designed a randomly moving bar paradigm (Fig. 3g, row #3). The bar moved only 56.7° in one direction on average, and then abruptly changed speed and direction. This was called the “randomly” moving bar experiment. Here, 26% neurons showed significant aVEVS, which was far greater than chance, though lesser than the systematic condition (Extended Data Fig. 14). The percentage of unidirectional, bidirectional, and untuned-stable cells were qualitatively similar to the systematic stimulus experiments. (Extended Data Fig. 11). Thus, the aVEVS cannot arise entirely from sequential movement of the bar. The retrospective latencies were also unaffected (Extended Data Fig. 9). To directly ascertain the effect of predictability on aVEVS, we separately analyzed the randomly moving bar data in the first 1-second after stimulus direction flip, and an equivalent subsample of data from later (Extended Data Fig. 11, see methods). aVEVS was similar in these two conditions. Further, aVEVS were not systematically biased by the angular movement speed of stimulus, nor did hippocampal firing encode stimulus speed beyond chance levels (Extended Data Fig. 11).

Recent studies have reported representational drift, i.e. slow remapping of place cells over several days^39^. We measured the activity of the same cells for more than one day, and measured changes in aVEVS without any changes in stimuli for the predictably moving, systematic bar of light. There was a large remapping of aVEVS across two days, evidenced by very low correlation between the tuning curves of the same neuron across two days (Fig. 3i). This was not due to difference in novelty, because rats had experienced this stimulus for at least one week.

Thus, unlike all or none changes in place cells that show complete remapping with large, but not small changes in visual cues^40^, aVEVS responses were largely invariant as measured by the correlation coefficient of the tuning curves (Fig. 3i). They showed gradually larger change in aVEVS with progressively greater changes in the visual cues, ranging from pattern, then color, then predictability but largest with the passage of time. These results were partly mediated by the change in preferred angle under different conditions. But, even when this contribution was factored out, a similar pattern of changes was observed (Extended Data Fig. 11).

### Most VEVS ne urons are place cells

During spatial exploration, majority of rodent hippocampal neurons show spatially selective responses, aka place cells. What is the relationship between aVEVS and spatial selectivity of neurons? We measured the activity of the same set of CA1 neurons, on the same day, during the aVEVS protocol and while rats freely foraged for randomly scattered rewards in two-dimensional environments (Fig 4a, see methods). 79% (184 out of 234) of neurons that were active in the bar of light experiment were also active during spatial exploration. This is far greater than the fraction (∼20%) of place cells that are active in two different environments during spatial exploration. Further, the firing rates during exploration and moving bar experiments were strongly correlated (Extended Data Fig. 15). Amongst cells that showed significant aVEVS, 90% (70 out of 78) showed significant spatial selectivity. Notably, the strength of tuning was also significantly correlated between these two experiments (Fig. 5c). Thus, despite very different experimental conditions and behavior the majority of aVEVS cells were also place cells, with similar activity and tuning.

**Fig. 4.**
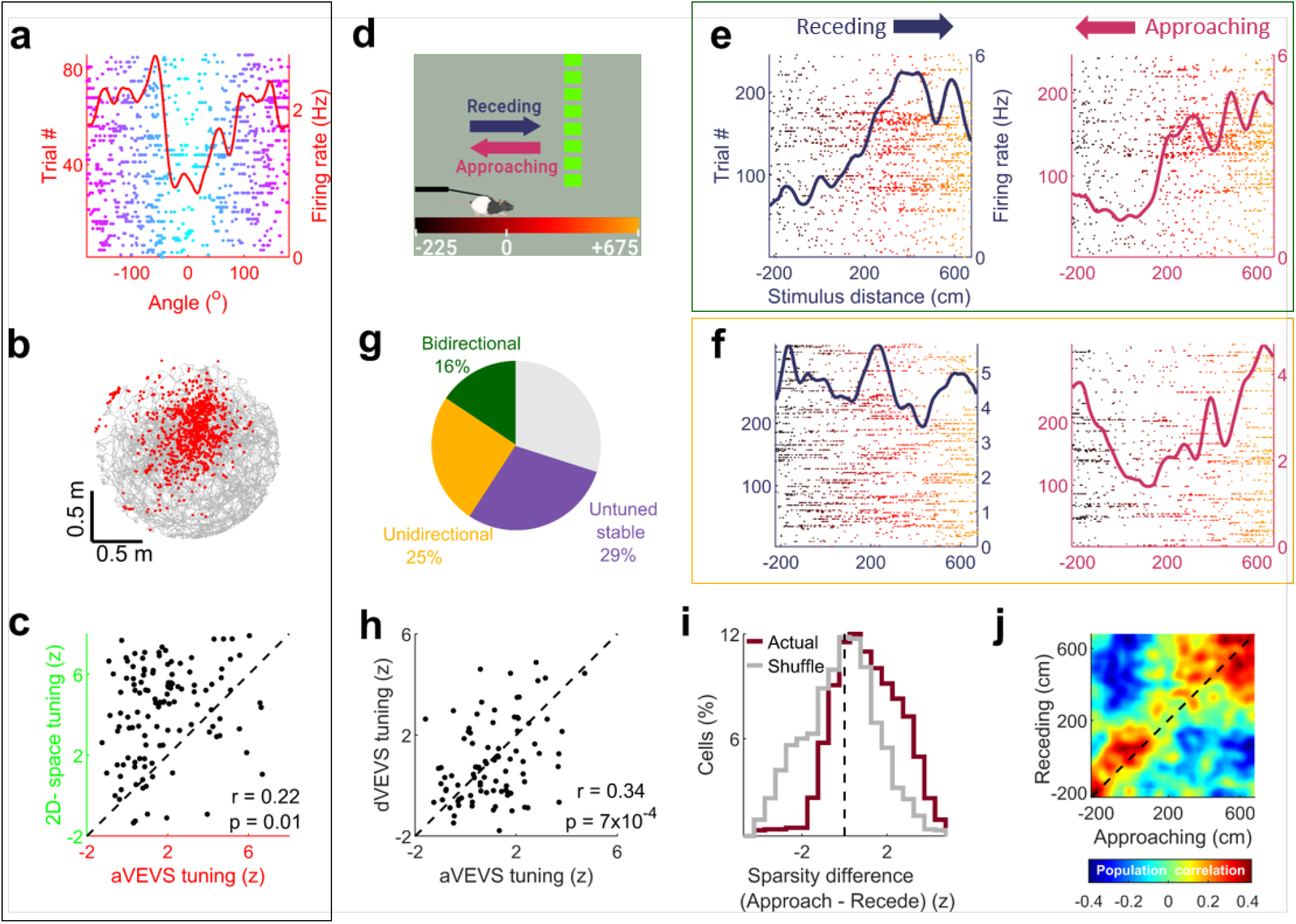
aVEVS cells are place cells and stimulus distance encoding cells. (**a)** A cell recorded on the same day showing significant aVEVS in the revolving bar of experiment and **(b)** allocentric spatial selectivity during free foraging on a circular table. Position of the rat (grey), spikes (red). **(c)** Strength of aVEVS and spatial selectivity measured by z-scored sparsity were significantly correlated (*Pearson* correlation indicated inset). **(d)** Schematic of the stimulus distance experiment. Bar of light moved towards and away from the rat at a fixed angle (0°). **(e)** Raster plots and firing rates of a bidirectional cell with significant tuning to the approaching (pink, top) as well as receding (dark blue, bottom) bar of light. Trial number (y-axis on the left) and firing rates (y-axis on the right) **(f)** Same as (e), but for a unidirectional cell, tuned for stimulus distance only during the approaching stimulus movement. **(g)** Relative percentages of cells, similar to. Fig. 2d. **(h)** Angular and linear stimulus position tuning was positively correlated (*Pearson* correlation indicated inset). **(i)** Stimulus distance tuning is larger during approaching epochs, even after down sampling spikes to have same firing rate (KS-test actual *p*=4.6×10^−4^, shuffle *p=*0.06). **(j)** Population vector overlap between responses in approaching and receding stimulus movement shows retrospective response, with maxima at values above the diagonal, similar to Fig. 3h, corresponding to a median latency of 63.8cm or 181.1ms (±377.6ms).

Spatial exploration involves not only angular optic flow but looming signals too. Hence, in the same apparatus, we measured 147 cells when the stimulus moved towards or away from rat, at a fixed angle of 0°, completing one lap in 10 seconds (Fig 5d-f). Similar to the revolving bar experiments, the animal was body-restricted, and his movements had no impact on the motion of the bar or rewards (Figure 1). The firing rates of 41% of neurons showed significant modulation as a function of the stimulus distance, i.e., dVEVS (Fig. 5g) and 27% of cells had untuned but stable responses. Neurons not only encoded distance but also direction of movement, with 17% and 8% of neurons showing significant tuning to only the approaching (moving towards) or receding (moving away) bar of light, respectively. For cells recorded in both stimulus distance and angle experiments (see methods), firing rates (Extended Data Fig. 12) as well as the strength of tuning were correlated, suggesting that the same population of neurons can encode both distance and angle (Fig. 5h). The preferred distance (i.e., the position of maximal firing) for the bidirectional cells was not uniform but bimodal, with over-representation near the rat (0 cm) or farthest away locations (500cm) (Extended Data Fig. 12). Neural firing rates were quite similar for approaching and receding stimuli, but stimulus distance coding was much stronger for approaching movements (Fig 5i). Retrospective response was also seen in dVEVS (Fig. 5j), with the population overlap between approaching and receding responses shifted to values above the diagonal (Fig. 5j). This corresponds to a retrospective shift of 70.6cm or 196.1ms.

## Discussion

These results demonstrate that a moving bar of light can reliably generate selectivity to distance, angle and direction of motion in hippocampal place cells, without any task demand, memory, reward contingency or locomotion requirements. Selectivity to these three spatial variables demonstrates visually evoked vectorial selectivity i.e., VEVS; unlikely to arise due to non-specific variables (Extended Data Fig. 13 for reward related controls, Fig. 14 for behavior related controls, Fig. 15 for GLM estimates, Fig. 16 for simultaneously recorded neurons showing diverse aVEVS and Fig. 17 for quantification of co-fluctuation of firing responses). Similar to place cells, only a few hundred aVEVS neurons were sufficient to accurately decode the angular position of stimulus. Positions in front of the rat and near him were overrepresented, similar to the visual cortices. Majority of neurons that encoded the bar position were also spatially selective during real world exploration and the strength of VEVS and spatial tuning were correlated. However, unlike place cells that shut down completely outside the place field, VEVS showed significant activity outside the preferred zone. While, place cells remap when the behavior is sequential vs random^38^, the aVEVS was relatively unchanged when the predictability or sequential nature of stimuli was altered.

Strikingly, while VEVS was retrospective, hippocampal place cells during spatial exploration ^5–8,41^ and head direction cells in related areas^42–44^, including in virtual reality setup similar to that used here^8^, show prospective or predictive responses (Extended Data Fig. 8). Thus, self-motion signals maybe required to turn the retrospective VEVS into prospective ones, necessary for navigational maps^12^. Indeed, robust responses and prospective coding were seen in virtual reality, but for relative distance, not absolute position, since only the optic flow and locomotion cues were correlated at identical distance^8^.

These results show that during passive viewing, rodent hippocampal activity patterns fit the visual hierarchy^45^. For example, the aVEVS show similar but smaller nasal-temporal magnification as visual cortex, e.g. larger tuning curve width for more peripheral stimuli, and over-representation of the nasal compared to temporal positions^27^. aVEVS is weaker but not absent for stimulus locations behind the rat, suggesting history dependence or other downstream processing providing stimulus information when it is not directly visible. History-dependence could also explain the unidirectional responses in our experiments, also seen in the primary visual cortex^46^, perhaps arising from similar plasticity mechanisms^47^. Further, like the visual cortex, hippocampal neurons too showed retrospective responses, but with larger response latency, suggesting visual cortical inputs reached hippocampus to generate VEVS. The larger latency is remarkably similar to that in the human hippocampus^48^, and in the rodent cortico-entorhinal-hippocampal circuit during Up-Down states^49–51^. However, there were no off responses in the VEVS and the tuning curves were broader and more unidirectional than in the primary visual cortex. This could arise due to processing in the cortico-hippocampal circuit, especially the entorhinal cortex^51^, or due to the contribution of alternate pathways from the retina to the hippocampus^52^.

Hippocampal spatial maps are thought to rely on the visual cues^3^. Rats can not only navigate using only vision in virtual reality, but they preferentially rely on vision^26^. Further, rats can navigate in virtual reality without robust vestibular cues, and neurons show robust selectivity to distance traveled and direction of hidden reward location, that predicts behavioral performance^12^. Consistent with the multisensory pairing hypothesis^9,30^, these multiplexed responses could arise when visual cue evoked VEVS are paired with locomotion and reward cues during the navigation task. In the absence of any correlation between multisensory stimuli, hippocampal neurons generate invariant, non-abstract and retrospective VEVS which are less robust than place cells. In the real world navigation tasks, the greatly enhanced correlations between all the internal and external multisensory cues could be encoded more robustly via Hebbian plasticity to generate anticipatory or prospective coding of absolute position^12,28,53^. Thus, the hippocampus can be driven reliably by a moving visual cue, similar to visual cortex, to generate a vectorial representation of space in a polar coordinate frame centered on the rat’s head. When combined with self-motion, rewards and multisensory cues, this could elicit not only allocentric place^2^ cells but task related hippocampal complexity^12^.

## Acknowledgements

We thank K. Delao, C. Polizu and S. Samant for help with data collection; V. Yuan, S. Ryklansky, A. Chorbajian and W. Zhu for help with single unit clustering; and D. Dixit for help with illustrations. This work was supported by grants to M.R.M. from the W.M. Keck Foundation, AT&T, NSF 1550678 and NIH 1U01MH115746.

## Author contributions

M.R.M. and C.S.P. designed the experiments. S.D., C.S.P., R.R., C.V., T.T., A.H. and K.C. performed the experiments. C.S.P. developed the stimuli and performed the analyses with input from M.R.M. M.R.M. and C.S.P wrote the manuscript with critical input from S.D. and other authors. **Competing interests:** The authors declare no competing interests.

## METHODS

All experimental procedures were approved by the UCLA Chancellor’s Animal Research Committee and were conducted in accordance with USA federal guidelines.

### Statistical tests

All usages of the Kolmogorov Smirnov test were with 2-sample, 2-sided hypothesis testing. All Student’s t-tests used here were one sample and used to test if the distribution was centered at zero. Unless otherwise noted, correlation coefficients were computed as Pearson coefficients using the *corrcoef* function in MATLAB. Circular tests used to compare angular quantities in Figure 3 and Extended Data Fig 7 and 11 were executed with the *circ_stats* toolbox in MATLAB^54^. Owing to the lack of multiple comparisons, no adjustments were necessary, except for the use of CCW as well as CW responses of the same cell in considerations of significantly tuned population.

### Subjects

Eight adult male Long-Evans rats (3 months old at the start of experiments) were individually housed on a 12-hour light/dark cycle. Their total food intake (15-20 g of food per day) and water intake (25-35 ml of water per day) were controlled and monitored to maintain body weight. Rats received 10-12ml of water in a 20-minute experiment. All experimental procedures were approved by the UCLA Chancellor’s Animal Research Committee and were conducted in accordance with USA federal guidelines.

### Experimental apparatus

Rats were body restricted with a fabric harness as they ran on an air-levitated spherical treadmill of 30cm radius. The rat was placed at the center of a cylindrical screen of radius 33cm and 74 cm high. Visual cues were projected on the screen. Although the rat was free to run and stop voluntarily, his running activity was decoupled from the projector and hence had no effect on the visual cues. Body restriction allowed the rat to scan his surroundings with neck movements. Running speed was measured by optical mice recording rotations of the spherical treadmill at 60Hz. Head movement with respect to the harnessed and fixed body was recorded at 60Hz using an overhead camera tracking two red LEDs attached to the cranial implant using the methods described before^8^. Rewards were delivered at random intervals (16.2 sec ±7.5s, 2 rewards, 200ms apart) to keep the rats motivated and the experimental conditions similar to typical place cell experiments.

### Behavioral pre -training

All experiments were conducted in acoustically-and EMF-shielded rooms. The rats were conditioned to associate a tone with sugar-water reward. They were gently body-fixed in the apparatus that allowed them to move their heads with respect to the body, but the body could not turn around. In order for the rats to remain calm in the apparatus for long periods, they were trained to navigate in a visually rich virtual maze where a suspended, striped pillar indicated rewarded position. After surgery, rats were exposed to the revolving bar environment for the first time, where the movement of the rat had no impact on the movement of the revolving bar. Six out of eight rats never experienced virtual reality after the revolving bar experiments began.

### Experiment Design

The salient visual stimulus was a 13 degrees wide vertical bar of light which revolved around the rat at a constant speed (10s per revolution) without any change in shape or size (Fig 1A). We used three different textures of visual cues as shown in Fig. 4. The results were qualitatively similar for all of them hence the data were combined. Each block of trials consisted of four clockwise (CW) or four counterclockwise (CCW) revolutions of the bar of light. There were 13-15 blocks of trials in each session. During the random bar of light experiment, the bar revolved at one of the six speeds: ±36°, ±72°, or ±108° per second and spanning angles ranging 30° to 70° at any given speed, before changing the speed at random. Reward dispensing was similar to the systematic bar of light experiment, with no relation to the angular position or speed of the stimulus. Manipulations of stimulus color, pattern, movement predictability and linearly moving stimulus were performed in a pseudo-random order in the same VR apparatus. Real world two-dimensional random foraging experiments and stimulus angle experiments were performed in a pseudo-random order, with an intermittent baseline of 25-40 minutes.

### Surgery

All rats were implanted with 25-30g custom-built hyperdrives containing up to 22 independently adjustable tetrodes (13μm nichrome wires) positioned bilaterally over dorsal CA1 (−3.2 to -4.0mm A.P., ±1.75 to ±3.1mm M.L. relative to Bregma). Surgery was performed under isoflurane anesthesia and heart rate, breathing rate, and body temperature were continuously monitored. Two ∼2 mm-diameter craniotomies were drilled using custom software and a CNC device with a precision of 25μm in all 3 dimensions. Dura mater was manually removed and the hyperdrive was lowered until the cannulas were ∼100 μm above the surface of the neocortex. The implant was anchored to the skull with 7-9 skull screws and dental cement. The occipital skull screws were used as ground for recording. Rats were administered ∼5mg/kg carprofen (Rimadyl bacon-flavored pellets) one day prior to surgery and for at least 10 days during recovery.

### Electrophysiology

The tetrodes were lowered gradually after surgery into the CA1 hippocampal sub region. Positioning of the electrodes in CA1 was confirmed through the presence of sharp-wave ripples during recordings. Signals from each tetrode were acquired by one of three 36-channel head stages, digitized at 40 kHz, band pass-filtered between 0.1Hz and 9 kHz, and recorded continuously.

### Spike sorting

Spikes were detected offline using a nonlinear energy operator threshold, after application of a non-causal fourth order Butterworth band pass filter (600-6000Hz). After detection, 1.5ms spike waveforms were extracted. A custom built “PyClust” software was used for spike sorting as described before ^8^.

### Tuning curve s and *z-*score calculation

Procedures similar to that described previously were used^8^. We binned the angular occupancy of the vertical bar and spikes in *N=120* bins of width 3° each and smoothed it with a Gaussian of σ =12°. Clockwise and counterclockwise movement directions were treated separately. To quantify the degree of modulation we computed sparsity *s* of an angular rate map where *r*_*n*_ is the firing rate in the 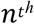 angular bin:

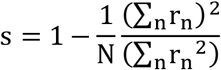

To assess the statistical significance of sparsity, we used a bootstrapping procedure, which does not assume a normal distribution. Briefly, for each cell, in each movement direction, spike trains as a function of the vertical bar from each block of trials were circularly shifted by different angles and the sparsity of the randomized data computed. This procedure was repeated 250 times with different sets of random value shifts. The mean value and standard deviation of the sparsity of randomized data was used to compute the z-scored sparsity of actual data using the function *zscore* in MATLAB. The observed sparsity was considered statistically significant if the z-scored sparsity of the observed spike train was greater 2, which corresponds to *p* < 0.0228 in a one sided t-test.

Similar procedure was employed for testing the significance angular tuning in the random bar of light condition. To keep the analysis comparable to systematic condition, spike trains were circularly shifted with respect to behavioral data by different random amounts for each block of 40 seconds, which is comparable to the time taken by the systematic visual cue to undergo four revolutions.

In addition to sparsity, we quantified aVEVS using several other measures.

Angle Selectivity index

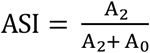

where A_2_ is the second harmonic component from the Fourier transform of the binned aVEVS response and A_0_is the DC level. This formulation of ASI is analogous to Orientation selectivity index (OSI), which is widely used in visual cortical selectivity quantification^54–56^

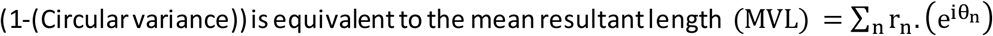

Where *r*_*n*_ is the firing rate in the 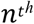 angular bin θ_n_ is the angular position corresponding to this bin and n is summed over 120 bins.

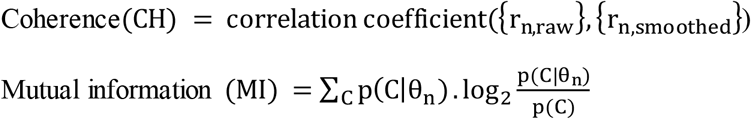

Where p(C) = ∑_n_p(θ_n_). p(C|θ_n_)

and C is the average spike count in 0.083 second window which corresponds to 1 angular bin that is 3° wide. Statistical significance of these alternative measures of selectivity was computed similar to that for sparsity and is detailed in Extended Data Fig. 1.

### Tuning curve width quantification

Full width at quarter maxima of the aVEVS rate map was computed around the maxima of the firing rate, i.e., the preferred angle, as the width at which the tuning curve first crossed 0.25 times the peak value. We chose 0.25 of maximum and not 0.5, i.e., FWHM as commonly done, because the tuning curves are often very broad with nonzero activity at nearly all angles, which is missed by FWHM.

### Modulation Inde x calculation

Firing rate modulation index of stimulus angle tuning (used in Fig. 1g) was quantified as *(FR*_*within*_*-FR*_*outside*_*) / (FR*_*within*_ *+ FR*_*outside*_), where *FR*_*within*_ and *FR*_*outside*_ are average firing rates in their respective zones. Similar definition of FR modulation index was used in Fig. 2g, to quantify the effect of uni-directional tuning inside and outside of the preferred zone, as *(FR*_*tuned*_ *- FR*_*untuned*_*) / (FR*_*tuned*_ *+ FR*_*untuned*_), where *FR*_*tuned*_ and *FR*_*unturned*_ are the average firing rates in the respective directions. Similarly in Fig. 4k, to quantify the effect of stimulus speed, as *(FR*_*fast*_ *– FR*_*slow*_*) / (FR*_*fast*_ *+ FR*_*slow*_), where *FR*_*fast*_ and *FR*_*slow*_ are the average firing rates during stationary epochs of respective stimulus movement speeds.

### Spike Train thinning

Neurons with larger number of spikes, e.g., due to longer experiments, have greater sparsity than when the number of spikes is less. To remove this artifact and compare the degree of aVEVS across all neurons and conditions, we employed a spike thinning procedure. Randomly chosen spikes were removed such that that the effective firing rate became 0.5 Hz for all neurons and then computed the sparsity of this thinned spike train (Extended Data Fig. 6). This procedure was used separately for CW and CCW directions to allow comparison of the degree of tuning in both directions, independent of the firing rate changes.

### Stability Analysis

Stability of neural angular tuning was quantified for CW and CCW directions separately. All the trials were split into two randomly chosen, equal and non-overlapping groups (∼30 trials each) and separate tuning curves computed for each half, with 120 equally spaced, non-overlapping angular bins. The correlation coefficient was computed between these two groups (*C*_*actual*_), which is a measure of stability. To compute the significance of stability, this procedure was repeated 30 times, with different random grouping of trials, and correlation coefficient computed between the two groups computed each time. This provided a distribution of thirty values of stability *C*_*actual*_. Same procedure was used for rate maps computed using random data (see z-score methods above) and correlation computed between two groups to obtain thirty different values of *C*_*random*_. A cell’s VEVS was considered significantly stable if the following conditions were met: the nonparametric rank-sum test comparing the thirty *C*_*actual*_ with thirty *C*_*random*_ was significant at *p*<0.05 and *C*_*actual*_ *> C*_*random*_. Untuned-stable responses were identified as responses with significant stability, but non-significant tuning (sparsity (*z*) < 2) and treated as a separate population in Fig. 2.

### Population Vector Overlap

To evaluate the properties of a population of cells, sessions were divided into trials in the CCW and CW movement directions of the visual bar. Population vector overlap between CCW and CW movement direction at angles (*⊖*_*r*_, *⊖*_*m*_) for N single units was defined as the Pearson correlation coefficient between vectors *(µ*_*1,r*_, *µ*_*2,r*_, *… µ*_*N,r*_ *) & (µ*_*1,m*_, *µ*_*2,m*_, *… µ*_*N,m*_ ) where µ_i,p_ is the normalized firing rate of the *i*^*th*^ neuron at *p*^*th*^ angular bin. Correlation coefficient of these sub-populations taken across angles indicates the existence of retrospective coding (Fig. 3f, Extended Data Fig. 9 and Fig. 4j). Similarly, for computing coherence in either direction, population vector overlap between two groups of trials of the same bar movement direction (as defined above, stability analysis methods) was computed separately for CCW and CW trials (Extended Data Fig. 9). Populations of tuned, untuned but stable and untuned-unstable cells were treated separately.

### Decoding analysis

Using the stability labels as obtained from above, recorded cells were divided into three populations: tuned (sparsity *z*>2), untuned (sparsity *z*<2) and stable, untuned and unstable. All the trials across all the cells within each population were separated into two groups: ten, randomly chosen trials were treated as the ‘observed trials’ and this data were decoded using the firing rate maps obtained from the remaining trials or the ‘look-up trials’. Commonly used population vector overlap method was used between the lookup and observed trials using a window of 250ms. Briefly, at each 250ms time point in the ‘observed data’, the correlation was computed between the observed population vector and the lookup population vectors at all angles. The circularly weighted average of angles, weighted by the (non-negative) correlations provided the decoded angle. The entire procedure was repeated 30 times for different sets of 10 trials. The error was computed as the circular difference between the decoded and actual angle at the observed time. Decoding of the stimulus distance (Extended data Fig. 12) was done similarly but by finding the distance corresponding to maximum correlation between “look-up” and “observed” data, since circular averaging is unavailable for linear distance close and away from the rat.

### Same ce ll ide ntification

Spike sorting was performed separately for each session using custom software^8^. Identified single units were algorithmically matched between sessions to enable same cell analysis (Fig 3 and 4). All the isolated cells in one session were compared with all the isolated cells in another session under investigation. Each putative unit pair was assigned a dissimilarity metric based on the Mahalanobis distance between their spike amplitudes, normalized by their mean amplitude. Dissimilarity numbers ranged from 2.5×10^−5^ to 17.2 across all combinations of units between two sessions. Putative matches were iteratively identified in an increasing order of dis-similarity, until this metric exceeded These putative matches were further vetted, using an error index defined on their average spike waveforms.

### Estimating the inde pendent contribution of he ad position, running speed and stimulus angle using GLM

To compute the independent contributions of head position, running speed and stimulus angle, we employed an GLM based estimation of firing, using the *glmfit* function in MATLAB, as described recently^30^. Head position and running speed were decoded in GLM using basis functions consisting of sinusoids. The log of running speed was used to ensure similar amount of data in each bin, and bins with zero speed were assigned an arbitrary, small value, which was on average equal to half the minimum non-zero running speed. Spike train and behavior data were downsampled to 100ms bins. The extreme one percentile of head position data and top one percentile of running speed data was excluded to remove the effects of outliers and ensure a good fit. CCW and CW tuning curves for stimulus angle were computed seperately. The statistical significance of the resulting tuning curves was estimated by computing sparsity and a bootstrapping method described above and used recently^30^.

### Quantification of population remapping

To compute the amount of remapping of firing rate, strength of tuning, preferred angle of firing and similarity between CCW and CW aVEVS we used the responses of the same cells recorded from different experimental conditions and defined remapping metrices as firing rate modulation index, difference between z-scored sparsity, circular distance between the angles corresponding to maximal firing, correlation coefficient between the firing rate profiles and the peak value and angular latency corresponding to the cross correlation between their tuning curves in the two conditions. This calculation was repeated 100 times using a random permutation to break the same cell pairing, to obtain a null distribution. The mean and standard deviation of this distribution was plotted in Fig. 3h-j and Extended Data Fig. 11, and compared with the actual value of the corresponding remapping metric.

### Quantification of trial to trial variability of aVEVS

Angular movement of the visual stimulus was separated into different trials starting and ending at 0°, which is the angular position in front of the rat. Mean firing rate in each trial was obtained by binning the spikes in that trial into 120 angular bins (3 degrees wide), and finding the average value of firing rates in each bin. Similarly, mean vector angle and mean vector length were obtained using *circ_r* and *circ_mean* functions of the Circular Statistics toolbox in MATLAB^54^ either by using all trials or only those trials when atleast 5 spikes were recorded (each trial was 10s long, yeilding 0.5Hz lowerbound on mean firing rate).

### Quantification of co-fluctuation for simultane uosly recorde d Avevs

To determine if the varibility was correlated across simultaneously recorded tuned cells, a co-fluctuation index for firing rate was defined for all cell pairs as the Spearman correlation between the trial-wise firing rate vectors of both cells. *Co-Fluctuation*_*FR*_*= spearman({F*_*1,k*_*},{F*_*2,k*_*})* where *F*_*i,k*_ denotes the mean firing rate of i^th^ cell on the k^th^ trial. Bootstrapping procedure to access significance of this index was employed by obtaining 100 shuffled indices when the order of trials was randomly reassigned. Similarly, to estimate the co-fluctation of aVEVS, we defined a similarity metric for each trial as S_i,k_ = crcf(*r*_*n,k*_, *R*_*n*_) where *n* denotes the angular bins, *R*_*n*_ overall tuning curve, and r_n,k_ is the firing rate in the n^th^ bin for the k^th^ trial and *crcf* is the correlation coefficient function. Co-fluctuation of tuning was defined analogously as Co-Fluctuation _aVEVS_ = spearman({S_1,k_},{S_2,k_}), and bootstrapped similarly as the firing rate co-fluctuation index. Analogously, co-fluctuation of depth of modulation, was quantified with correlation coefficient of mean vector lengths (MVL) as Co-Fluctuation_MVL_ = spearman*({L*_*1,k*_*},{L*_*2,k*_*})*, where *L*_*i,k*_ denotes the mean vector length across stimulus angles of i^th^ cell on the k^th^ trial. Correlated changes in aVEVS preferred angule were quantified by Co-Fluctuation_MVA_ = circular correlation*({A*_*1,k*_*},{A*_*2,k*_*})*, where *A*_*i,k*_ denotes the mean vector angle of spiking of i^th^ cell on the k^th^ trial.

## EXTENDE DATA

**Extended Data Fig. 1.**
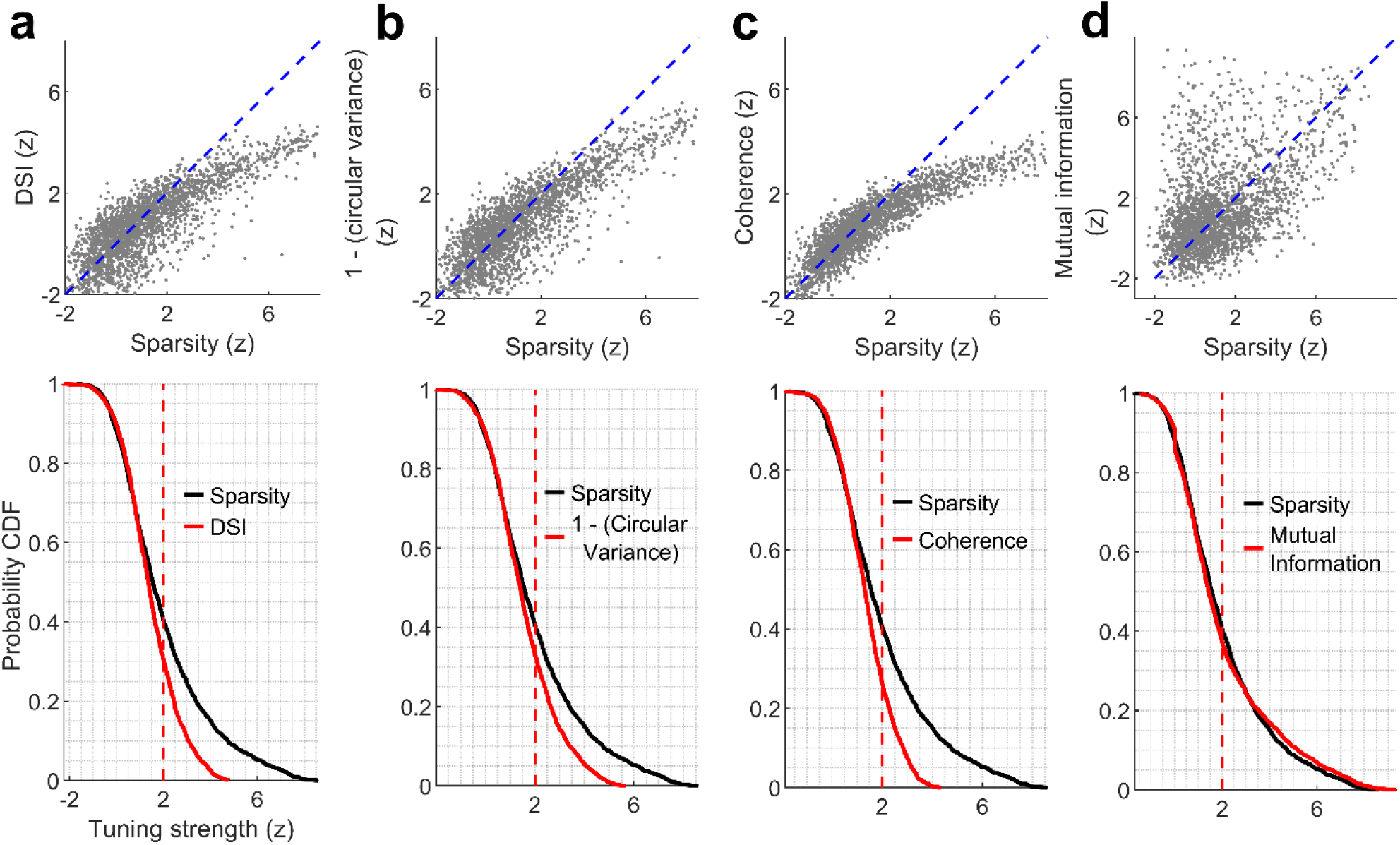
Relationship between different properties of aVEVS: **(a)** (top) aVEVS quantified by z-scored sparsity is significantly correlated (*r*=0.82, *p*<10^−150^) with, but significantly greater than the z-scored direction selectivity index (DSI) (41% z>2 for sparsity vs 31% for DSI, KS-test p=9.3×10^−10^). (Bottom) Cumulative histogram (cdf) of z-scored metric of sparsity and DSI. **(b)** Similar as (a), (1-(circular variance)) is significantly correlated (*r*=0.84, *p*<10^−150^) but significantly weaker (33% *z*>2 for (1-circular variance)) than sparsity. (KS-test *p*=7×10^−6^). **(c)** Similar to (a) coherence is significantly correlated (*r*=0.89 *p*<10^−150^) but significantly weaker (26% z>2 for coherence KS-test *p=*6.3×10^− 16^) than sparsity. **(d)** Similar to (a), but mutual information is significantly correlated (*r*=0.47 p=8.6×10^−132^) but significantly smaller than sparsity (37% z>2 for mutual information, KS-test *p*=7.2×10^−5^).

**Extended Data Fig. 2.**
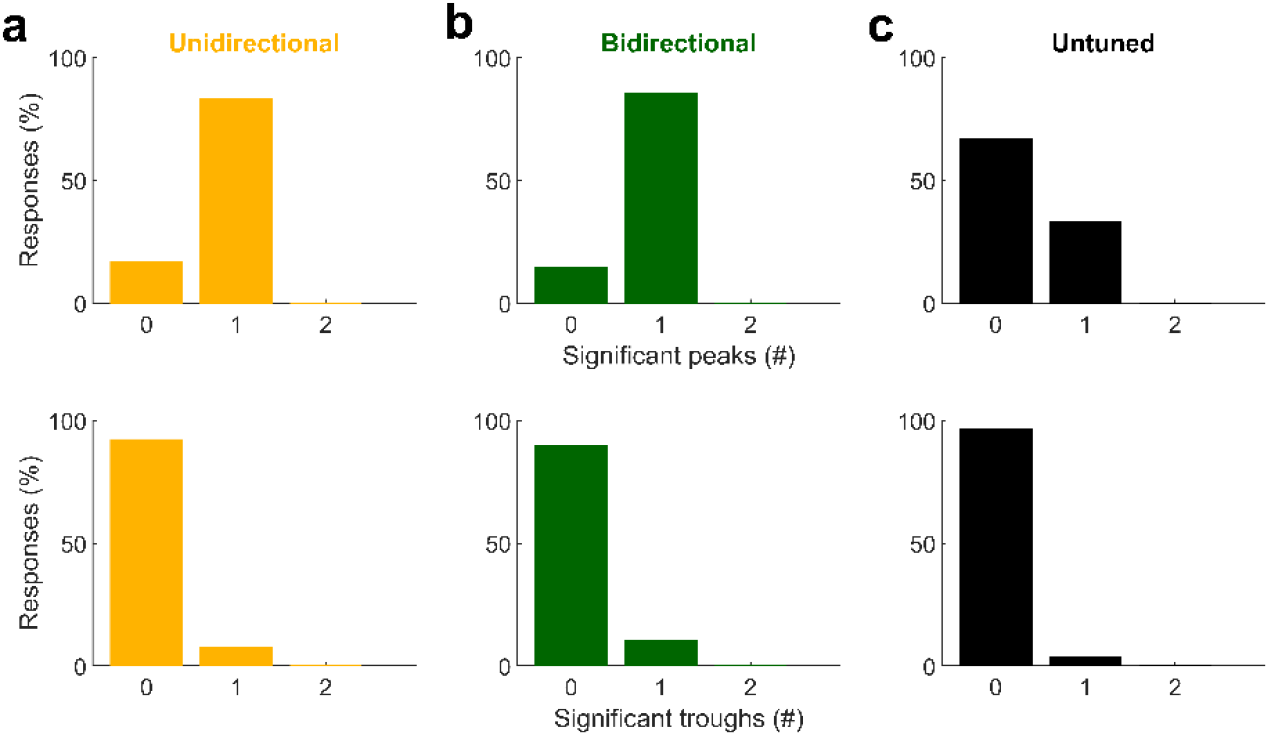
Unimodality of aVEVS: Majority of (**a**) uni-directional as well as (**b**)bi-directional tuning curves were unimodal with only one significant peak (top row), whereas (**c**) untuned responses did not have significant peaks, as expected. Both tuned responses were used for the bi-directional cells, and only the tuned response was used for the uni-directional cells. Significant troughs, i.e. off-responses were not found for unidirectional or bidirectional cells (bottom row). Significance of a peak (or trough) was determined with the spike train shuffling analysis, similar to that performed to compute the z-scored sparsity. A peak (trough) was determined to be significant if it was larger (smaller) than the median value of peaks (troughs) in all shuffles and had a height of at least 20% of the range of firing rate variation in the shuffle data. These criteria resulted in zero significant peaks for some tuned responses.

**Extended Data Fig. 3.**
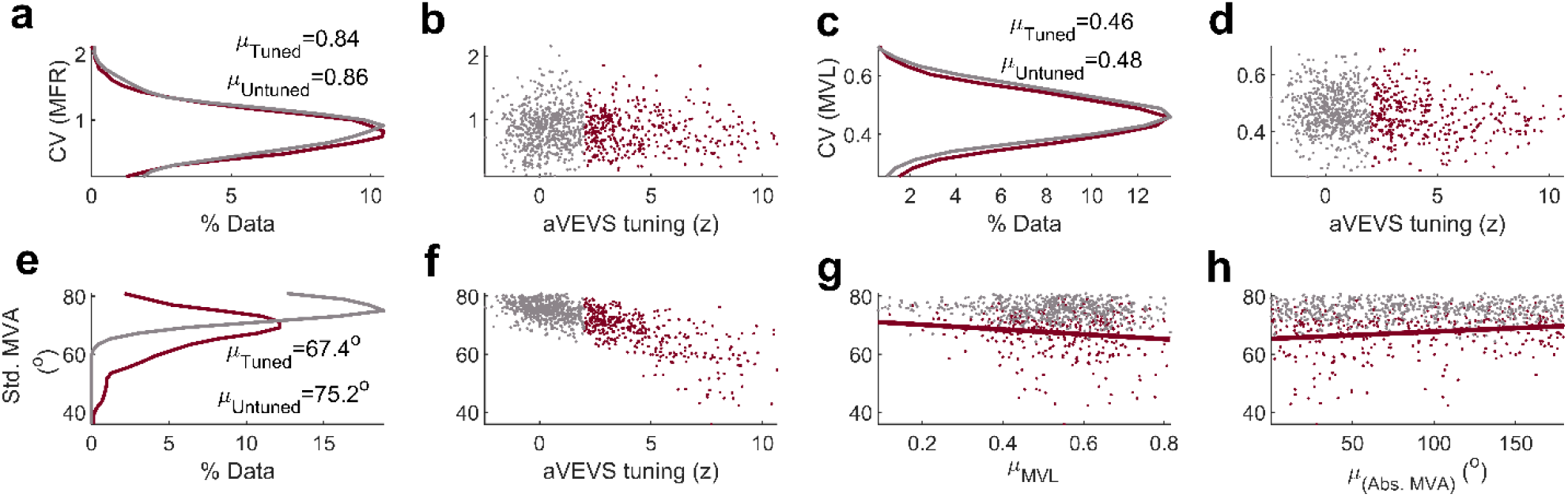
Trial-to-trial variability of mean vector angle but not mean firing rate determines aVEVS tuning. For each cell, in each trial, we computed the mean firing rate (MFR), mean vector length (MVL) and mean vector angle (MVA) of aVEVS (see *Methods*). To enable comparison across metrics, this analysis was restricted to responsive trials (firing rate above 0.5 Hz) where MVL and MVA could be meaningfully computed. Qualitatively similar results were obtained when this restriction was removed. **(a)** Trial to trial fluctuations of firing rate was qualitatively similar between tuned (maroon) and untuned (gray) cells (KS-test *p*=0.25). **(b)** The variability was not significantly correlated with the degree of aVEVS tuning (*Pearson* partial correlation, after factoring out mean firing rate, *p*=0.85). **(c)** The variance of MVL (see methods) was significantly greater for untuned cells (KS-test *p*=0.01) than tuned cells and **(d)** was inversely related to aVEVS tuning strength (*r=*-0.19, *p*=7.3×10^−10^). **(e)** The circular standard deviation of MVA, which signifies the instability of aVEVS tuning from trial to trial, was significantly (*p*=1.3×10^−72^) smaller (11%) for tuned than untuned cells and **(f)** strongly anti-correlated with aVEVS (*r*=-0.77 *p*=7.4×10^−192^). **(g)** This standard deviation of MVA was inversely correlated with MVL for tuned (*r*= -0.15 *p*=0.004), and for untuned cells (*r*= -0.12 *p*=0.003). **(h)** It was also positively correlated with the preferred angle of tuning (*r*=0.18 *p*=3.5×10^−4^), with lower variability for cells tuned to the front angles (0°) than behind (±180°). Standard deviation of MVA was uncorrelated with preferred angle of tuning for untuned cells (*r*=0.02, *p*=0.67). All correlations were computed as *Pearson* correlation coefficients.

**Extended Data Fig. 4.**
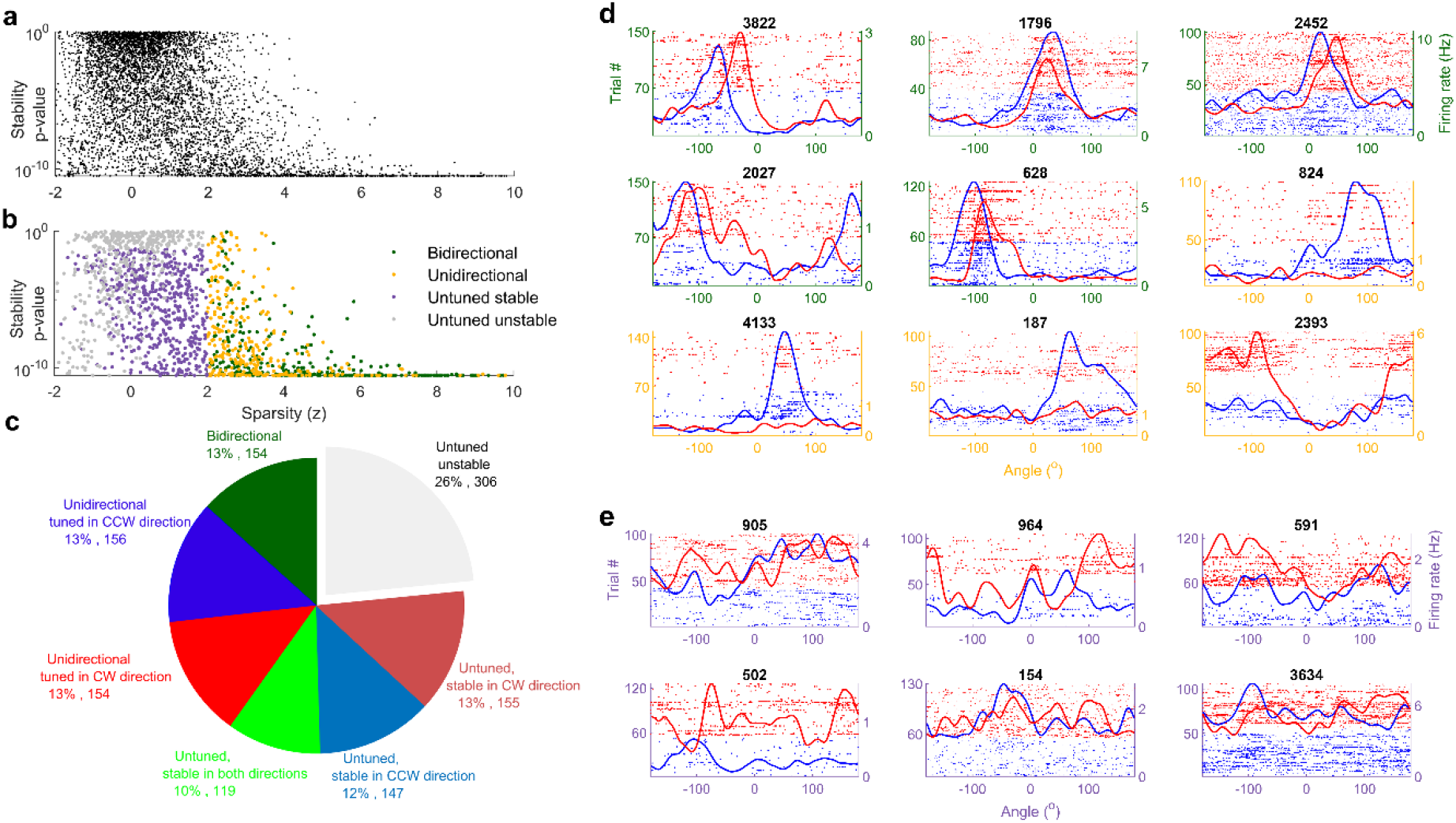
Continuity of stability and sparsity measures and example cells: across all neurons, the z-scored sparsity, i.e., degree of tuning, and stability varied continuously, with no clear boundary between tuned and untuned neurons. (**b**) Same distribution as (a), with color-coding of stable and tuned responses separated. (**c**) Detailed breakdown of aVEVS properties that had significant sparsity (i.e., tuned) or significant stability and whether these were observed in both directions (e.g., bidirectional stable) or only one direction (e.g. unidirectional tuned). If unidirectional, whether CW or CCW direction was significant. Nearly all cells that were significantly tuned in a given direction were also stable in that direction. **(d)** For clarity, the CCW (blue) and CW (red) trials are stacked separately in all raster plot figures, even though these alternated every four trials. First five examples are of bi-directionally tuned cells (green y-axis); next four examples are of uni-directionally tuned cells (orange-yellow y-axis). **(e)** These cells did not have significant sparsity (z<2) in either direction but had significant stability.

**Extended Data Fig. 5.**
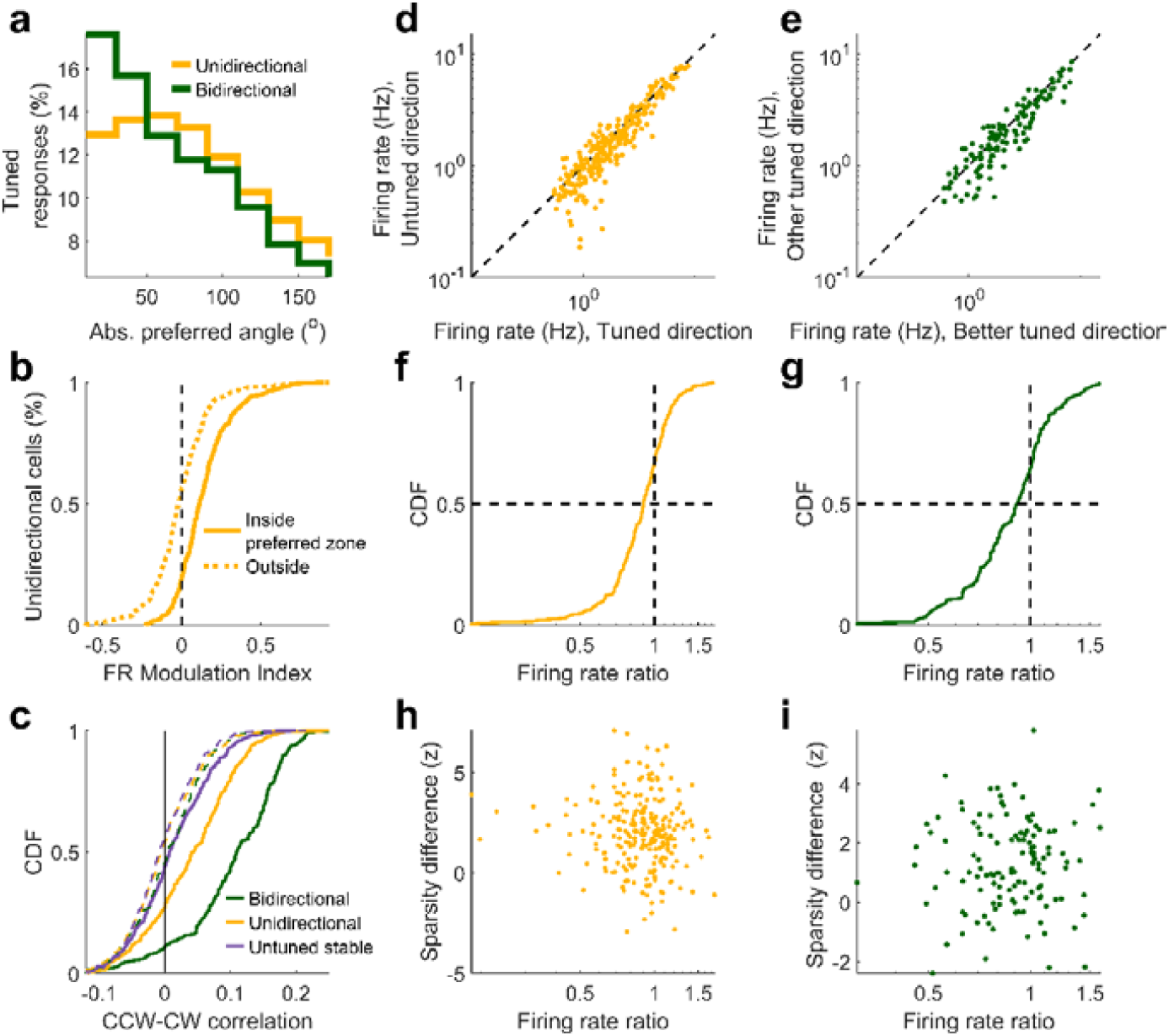
Firing rate differences between CW and CCW revolution direction: (**a**) Percentage of tuned responses as a function of the absolute preferred angle, for bidirectional and unidirectional populations are significantly different from each other (KS-test *p*=0.04).**(b)** Firing rate modulation index for uni-directional cells inside preferred zone was significantly different from zero (t-test, *p*=4.1×10^−35^), but not outside (*p*=0.35). **(c)** Correlation coefficient of CCW and CW responses for different populations of cells, (KS-test green, bidirectional, *p*=3.3×10^−27^, orange, unidirectional *p*=7.0×10^−27^, lavender, untuned stable, *p*=4.4×10^−4^). Dashed curves indicate respective shuffles. Firing rate of **(d**) unidirectional cells in tuned versus untuned directions shows significantly higher (KS-test *p*= 7.9 ×10^−9^) firing rates in the tuned direction **(e)** Same as (d), for bidirectional cells showing higher firing rate (KS-test, *p*=2.4×10^−18^) in the revolution direction with better tuning. **(f)** Cumulative histogram of ratio between firing rate in untuned to tuned direction was less than one for 67% (65%) of cells. **(g)** Same as (f), but for bidirectional cells (other/better since both directions are tuned) showing 65% of firing rate ratios were less than one. (**h**) To remove the contribution of firing rate to sparsity, the strength of tuning (z-score sparsity) difference was computed with spike thinning procedures (similar to Extended Data Fig. 8, see methods) ensuring equal firing rate in both directions. The difference in tuning strength (z-scored sparsity) was not significantly correlated with firing rate ratio for unidirectional (*r*=-0.09 *p*=0.16) as well as (**i**) bidirectional (*r*=0.005 *p*=0.95) populations. For bi-directionally tuned cells, aVEVS with higher z-scored sparsity was labeled as the “better” response, and the aVEVS with lower z-scored sparsity was called “other” response. All correlations were computed as *Pearson* correlation coefficients.

**Extended Data Fig. 6.**
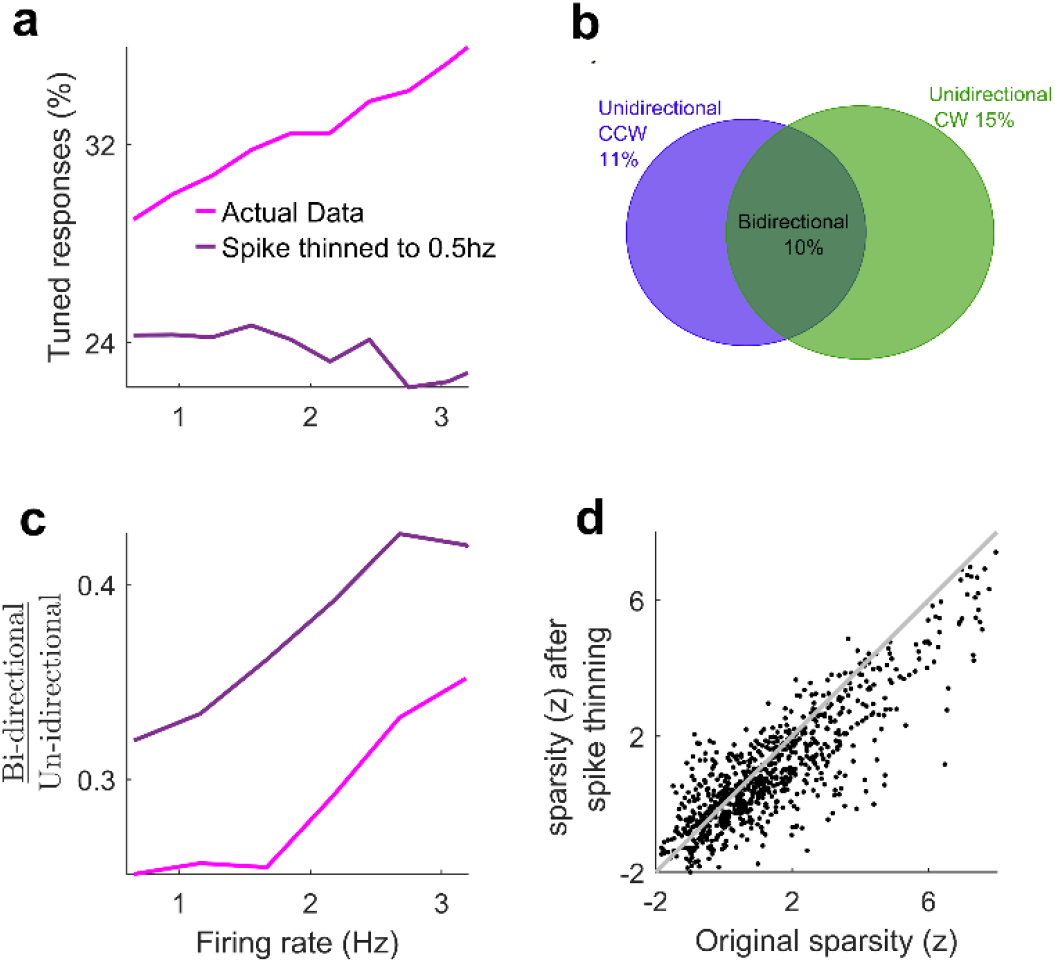
The relative number of bidirectional cells increases with mean firing rate, but not the fraction of tuned cells. To remove the effect of firing rate on z-scored sparsity computation, we randomly subsampled spike trains to have a firing rate of 0.5 Hz (see methods). (**a**) The fraction of cells with significant sparsity, i.e., fraction tuned, increased with the firing rate for the actual data (*r*=0.11 *p*=2.2×10^−6^), but after spike thinning, there was no correlation (*r*=0.01, *p*=0.77). Thus, the true probability of being tuned was independent of the firing rate of neurons. (**b**) Proportion of bidirectional and uni-directional tuned neurons is comparable (10% vs 13%) with and without spike thinning. (**c**) Fraction of bi-directional cells compared to uni-directional cells increases with original firing rate, even after spike train thinning. (**d**) Spike thinning procedure reduces the sparsity of the tuning curves, as expected, due to loss of signal. After spike thinning, sparsity was significantly correlated in both directions of revolution (*r*=0.39, *p*=3.8×10^−31^) and this was not due to the rate changes because sparsity was uncorrelated with firing rates (*r*=0.01, *p*=0.72 for CCW sparsity and firing rate, *r*=0.02, *p*=0.54 for CW sparsity and firing rate). All correlations were computed as *Pearson* correlation coefficients.

**Extended Data Fig. 7.**
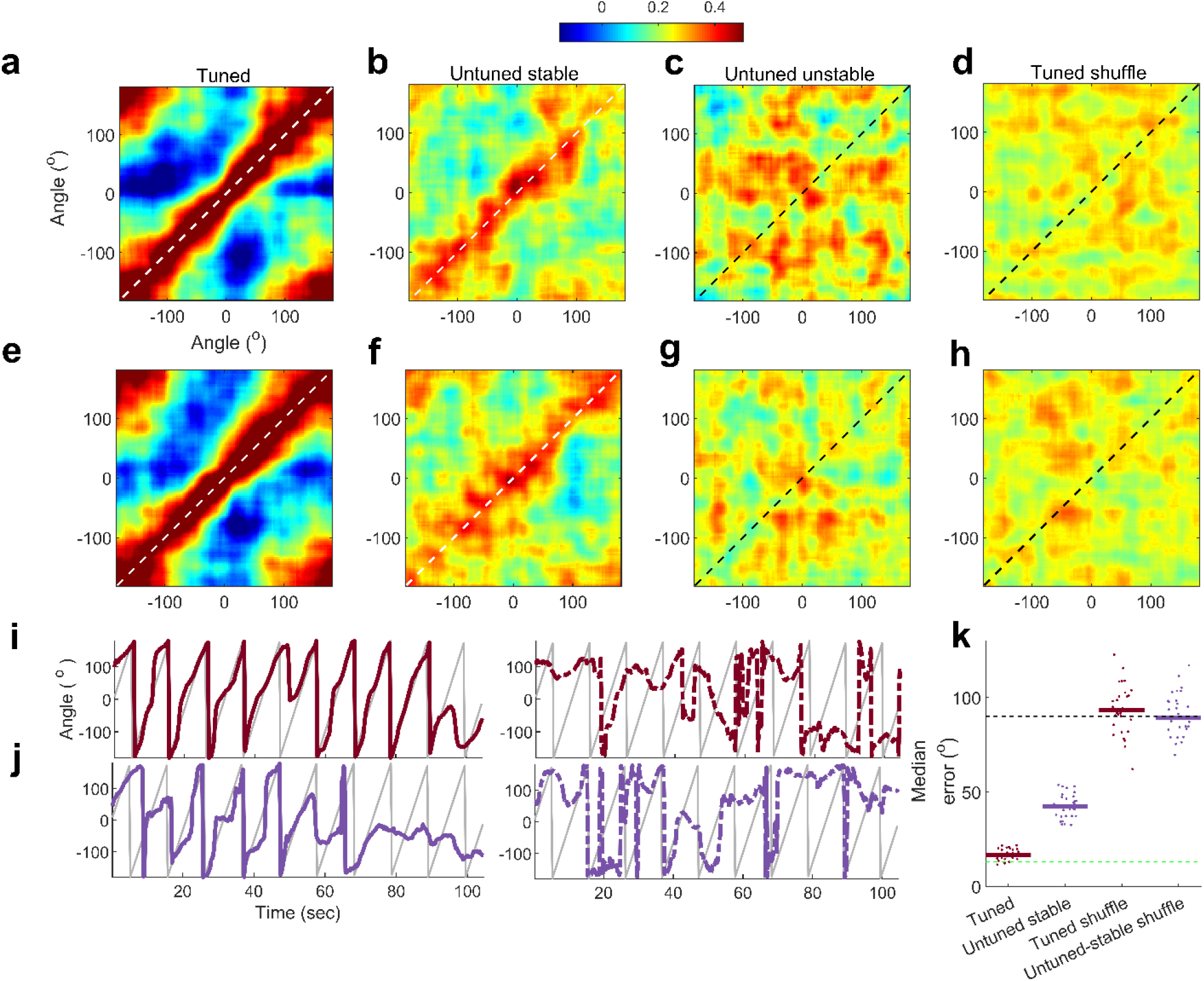
Population vector stability and decoding of visual cue angle. (**a**) Stability for CCW tuned responses (number of cells, n=310). Color indicates correlation coefficient between two non-overlapping groups of trials’ population responses (see methods). The maximum correlation values were pre-dominantly along the diagonal. Maxima along x-axis and y-axis were significantly correlated (Circular correlation coefficient *r*=0.97, *p*<10^−150^) (**b**) Same as (a) but using untuned stable cells (n=266) showed significant ensemble stability (*r*=0.91, *p*<10^−150^). (**c**) Same as (a) but using untuned and unstable cells (n=306). This was not significantly different than chance (*r*= -0.16, *p=*0.09). (**d**) Same as (a), using tuned cells with their spike trains circularly shifted in blocks of four trials, showed no significant stability (*r*=1.1×10^−3^, *p*=0.99). **(e)-(h)** Same as (a)-(d), but for CW data. **(i)** Decoding CW direction shows similar results as in CCW direction (shown earlier in Figure 2). Similar analysis as shown in Fig 2 for the stimulus movement in CW direction. (**Left**) Decoding cue angle in 10 trials of CW cue movement, using all other CW trials to build a population-encoding matrix. Gray trace indices movement of visual bar, colored trace is the decoded angle. **(Right)** Same as left, for shuffle data. (**j**) Same as (i) but using the untuned-stable cells in CW movement direction. (**k**) Median error between stimulus angle and decoded angle over 10 instantiations of decoding 10 trials each for actual and cell ID shuffle data. Green dashed line indicates width of the visual cue; black dashed line indicates median error expected by chance.

**Extended Data Fig. 8.**
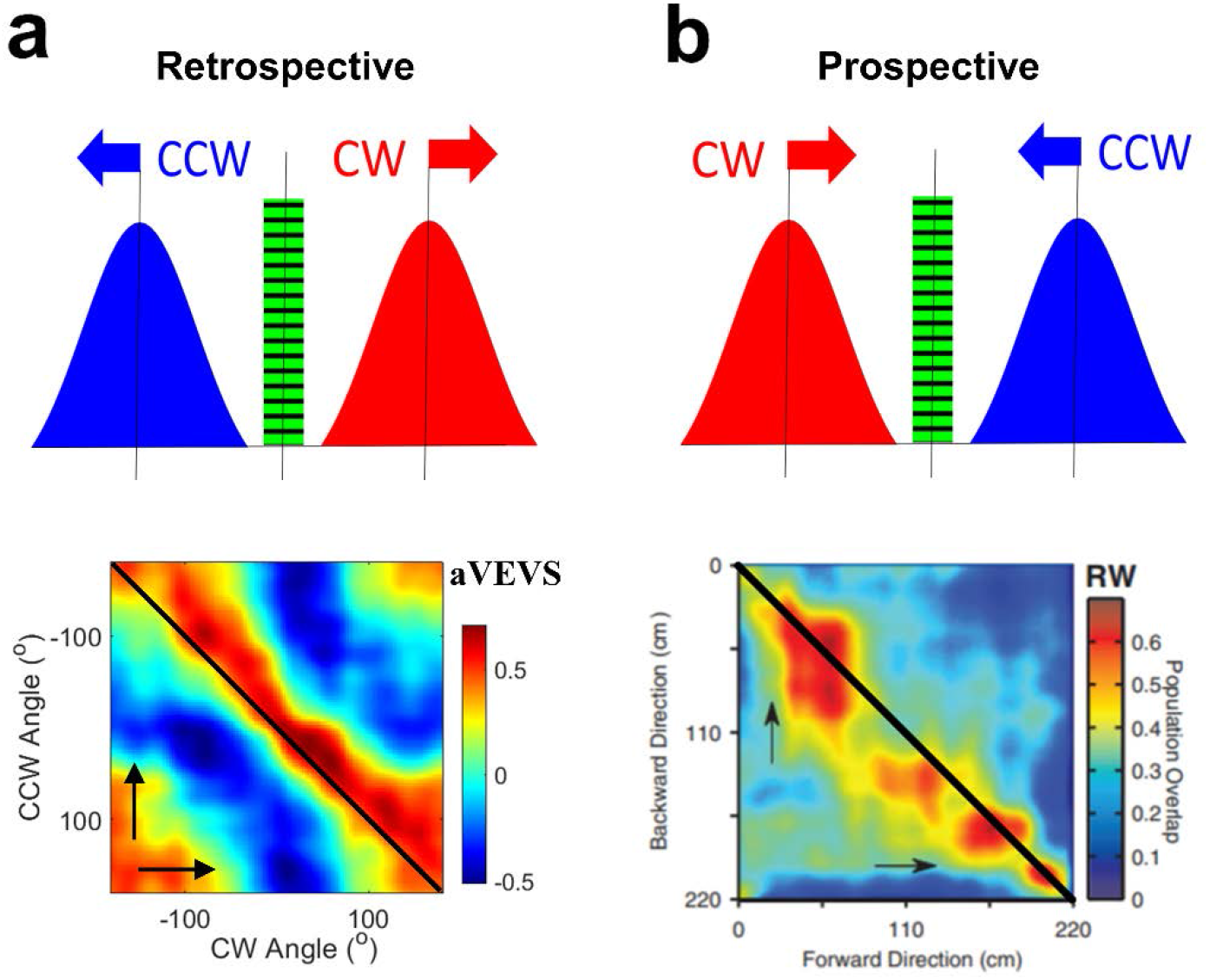
Retrospective coding of aVEVS cells versus prospective coding in place cells. **(a) Top-**A bidirectional cell responds with a latency after the stimulus goes past the angular position of the bar of light depicted by the green stripped bar. **Bottom**-Population overlap is above the 45° line, indicating retrospective response. **(b)** Same as (a) but for a prospective response, where the neuron responds before the stimulus arrives in the receptive field. Such prospective responses are seen in place fields during navigation in the real world, where the population overlap is maximal below the 45° line (adapted from earlier work^20^). Prospective coding was seen in purely visual virtual reality, but those cells encoded prospective distance, not position.

**Extended Data Fig. 9.**
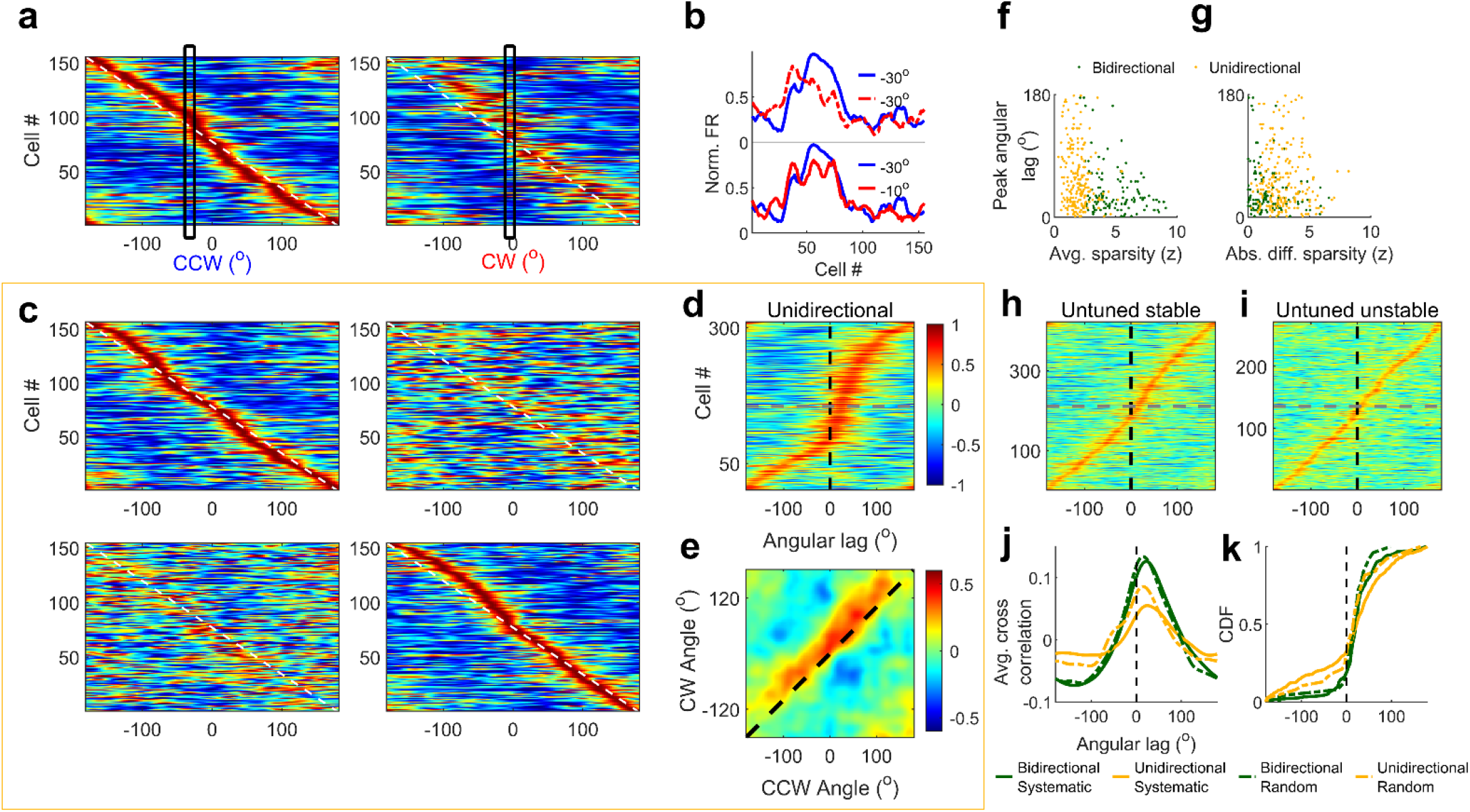
Significant retrospective aVEVS in the untuned stable cells but not unstable cells. (**a**) Stack plots of normalized population responses of cells, sorted according to the peak angle in the CCW (left). The corresponding responses of cells in the CW direction (right). (**b**) The firing rate, averaged across the entire ensemble of bidirectional cells at -30° in the CCW direction was misaligned with the same in CW direction at the same angle (top), but better aligned with the response at -10° (bottom, vertical boxed in (a)), showing retrospective response. (**c**) Same as (a) for uni-directional cells with CCW tuned cells (top row) and CW tuned cells (bottom row) sorted according to their aVEVS peak in the tuned direction. (**d**) Same as in (Fig. 3e) for unidirectional cells. Majority (67%) of the cross correlations between CCW and CW responses had a significantly positive lag (median latency=19.9°±86.1°, circular median t-test at 0°, *p*=1.8×10^−10^). The larger range of latencies and weaker correlations for unidirectional cells compared to the bidirectional cells could arise because significant tuning is present in only one direction. (**e**) Same as (Fig. 3f) for unidirectional cells. For all angles the population vector cross correlation coefficients had a peak at a positive lag (CW peak–CCW peak, median= +56.2° ±23.7° circular median t-test, *p=*1.5×10^−36^), which was not significantly different from the retrospective lag in bidirectional cells (KS-test, *p*=0.28). **(f)** Average strength of tuning in CCW and CW direction is inversely related to the peak angular lag between the two aVEVS for bidirectional (*Pearson’s r*=-0.19 *p*=0.04) as well as unidirectional cells (*Pearson’s r*=-0.16 *p*=0.02). **(g)** Absolute difference between strengths of tuning between CCW and CW directions was not significantly correlated with the peak angular lag in their cross correlation for bidirectional (*r*=0.13 *p*=0.14) or unidirectional cells (*r*=0.03 *p*=0.64). This analysis was restricted to cells with retrospective lags, which were in majority. (**h**) Untuned stable cells too show significant retrospective bias, quantified using the cross correlation between the tuning curves in CCW and CW directions (median lag =13.6° circular median t-test at 0°,*p*=0.02). (**i**) This is not seen for the untuned unstable population (median =4.6°, circular median t-test at 0°,*p*=0.39). (**j**) Cross-correlations between CCW and CW tuning curves were averaged across all the bidirectional cells (green curves) for the systematic (latency for peak=25.7°) and random (16.7°) condition and showed a similar pattern of retrospective coding. (two sample KS-Test to ascertain if the distribution of latencies was significantly different, *p*=0.75). Unidirectional cells showed similar pattern for systematic (19.7°) and random (31.8°) conditions, but correlations were weaker than bidirectional cells. (**k**) Cumulative distributions show that under systematic and random conditions comparable number of cells had positive latency (80% each) for bidirectional cells, and (67% and 68%) unidirectional cells respectively.

**Extended Data Fig. 10.**
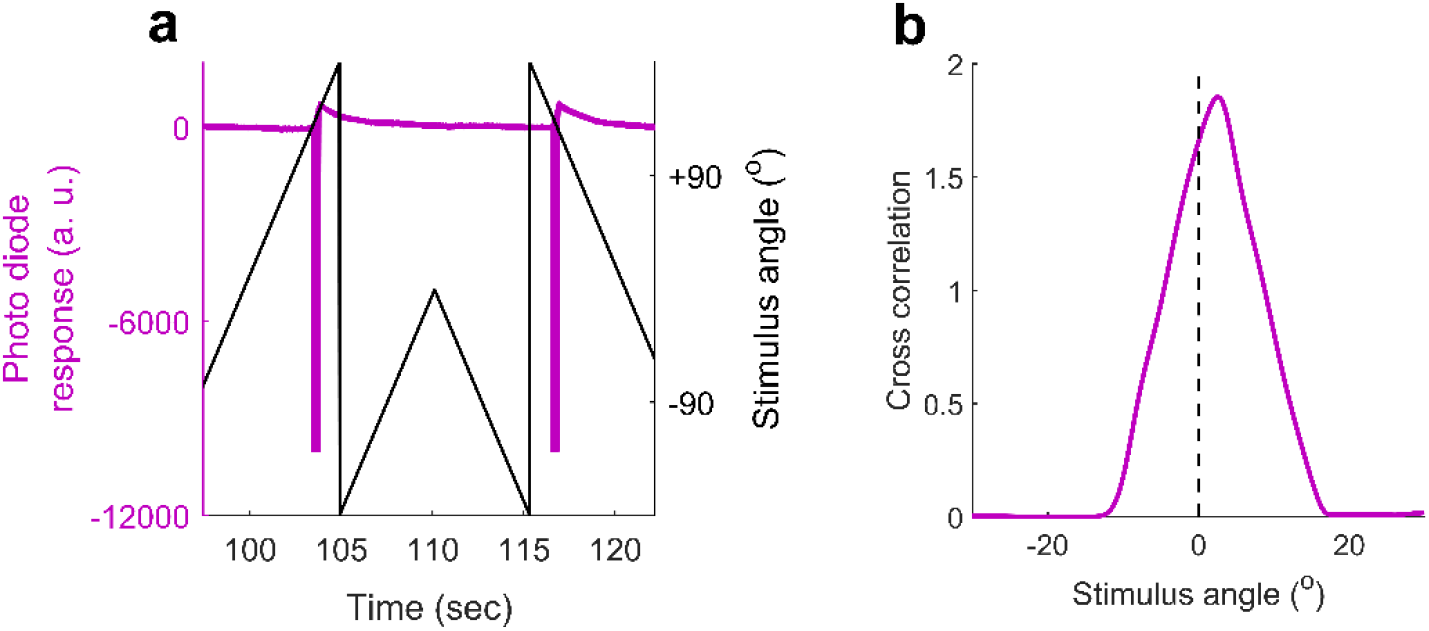
Photodiode experiment to measure the latency introduced by the equipment: Instead of a rat, we placed a photodiode where the rat sat. (**a**) The signal from the photodiode (purple trace) synchronized with bar position (black) was extracted and (**b**) cross correlation computed between the CW and CCW tuning curves of photodiode response. The cross correlation had maxima at a latency of -2.8°, which corresponds to a temporal lag of 38.9ms. This was much smaller than the latency between neural spike trains (median latency 22.7°, corresponding to a temporal latency of 315.3ms before accounting for the recording apparatus latency). For all the latency numbers reported in the main text, this small latency introduced by the recording apparatus was removed.

**Extended Data Fig. 11.**
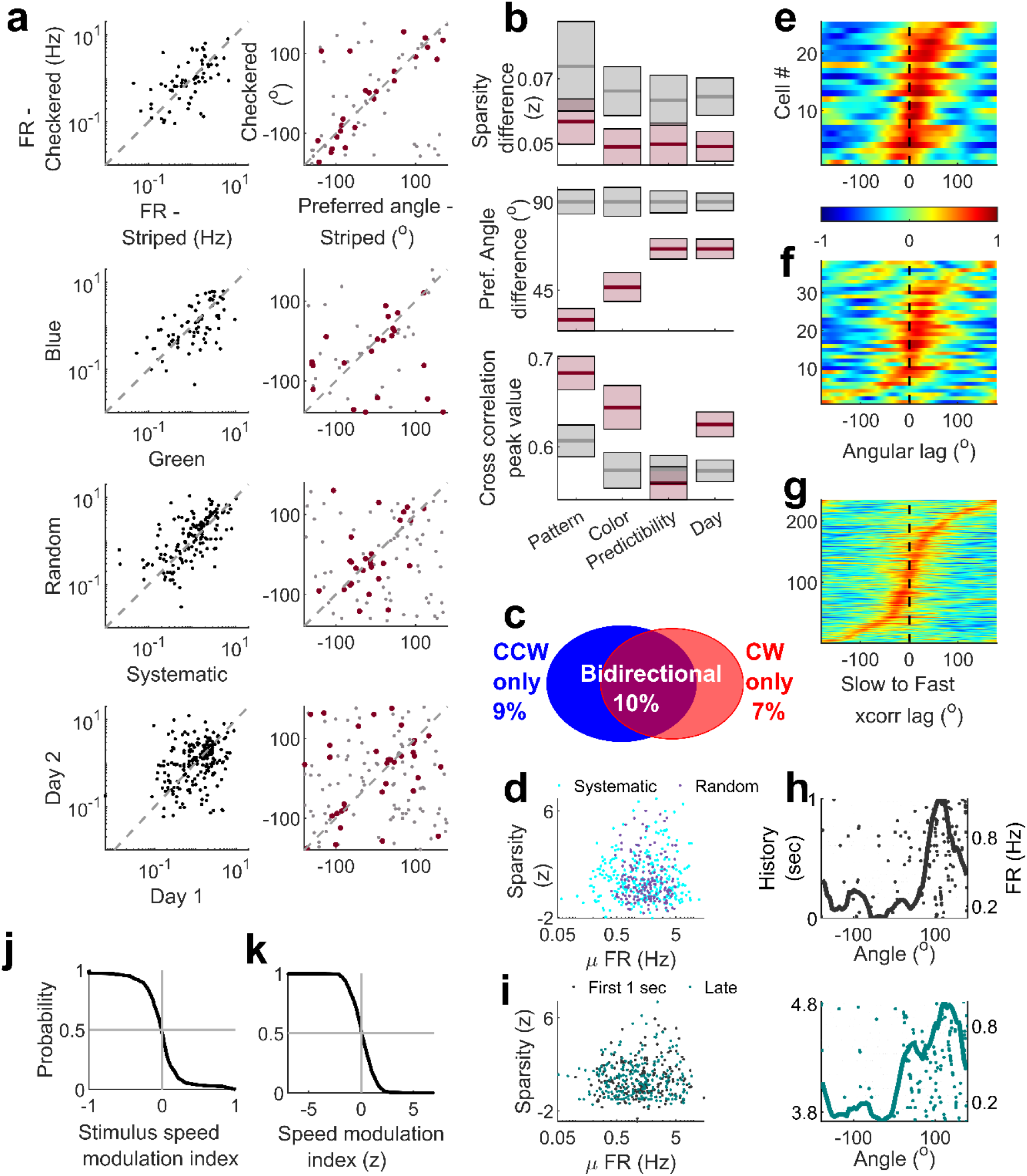
Additional properties of aVEVS invariance. **(a) (Row 1**) For same cells recorded in response to the movement of a green striped and green checkered bars of light, mean firing rate during stationary epochs (running speed< 5cm/sec), was significantly correlated (*Pearson’s r*=0.48 *p=* 2 × 10^−5^). Preferred angles of aVEVS between the two stimulus patterns were also significantly correlated (circular correlation coefficient, *r*=0.32 *p=*5×10^−3^). Solid red dots denote preferred angles of cells tuned (sparsity (z) > 2) in both conditions; gray dots are for cells with significant tuning in one of the conditions. **(Row 2)** Same as (a-Row 1), but for responses to changes of stimulus color, green and blue. Firing rate (r=0.45 *p=*1×10^−4^) & preferred angle (*r*=0.36 *p=*0.01) were correlated. **(Row 3)** Same as (a-Row 1), but for changes to predictability of the stimulus, also called “random” vs “systematic”. Firing rate (*r*=0.55 *p=*2×10^−13^) & preferred angle (*r*=0.27 *p=*0.01) were significantly correlated between systematic and random stimuli movement. **(Row 4)** Same as (a-Row 1), but for responses recorded across 2 days. Firing rate (*r*=0.28 *p=*3.2 × 10^−5^) & preferred angle (*r*=0.22 *p=*0.006) were correlated. **(b)** Similar to Fig 4, we computed the population remapping indices based on sparsity difference, preferred angle difference and peak value of cross correlation. The sparsity difference did not show a systematic pattern, but the other two metrics generally showed increasing remapping going from pattern (correlation=0.68, angle difference=30°) to color (correlation=0.64, angle difference=46.5°) to predictability (correlation=0.55, angle difference=66°) and across days (correlation=0.63, angle difference=66°). *n* indicates the number of responses measured in both conditions for each comparison, similar to Fig. 4e. Thick line – median, box – sem. **(c)** Percentage of tuned responses in the random stimulus experiments, showing, comparable bi-directionality (10% here vs 13% for systematically moving bar). **(d)** For same cells recorded in random and systematic stimulus experiments, the distributions of firing rates and selectivity, quantified by z-scored sparsity, were not significantly different (cyan-systematic, purple-random, KS-test for z-scored sparsity *p*=0.14, for firing rate *p*=0.27). **(e)** Cross correlation between CCW and CW tuning curves showing lagged response for the majority of bidirectional cells in the random condition. **(f)** Same as (e), but for unidirectional cells. **(g)** Cross correlation of tuning curves (for tuned cells in the random stimulus experiment) between fast- and slow-moving stimulus was calculated from the subsample of data for a particular speed in CW and CCW direction separately and stacked together after flipping the CCW data and was not significantly biased from zero (Circular median test at 0°, *p*=0.56). **(h)** Example cell showing similar aVEVS for data within 1 second of stimulus direction change(left), or an equivalent, late subsample(right). **(i)** Firing rate (KS-test *p*=0.73) and sparsity (KS-test *p*=0.87) were not significantly different for these two subsamples of experimental recordings. **(j)** In the randomly moving stimulus experiments, we computed a stimulus speed modulation index (see methods). This distribution was not significantly biased away from zero. **(k)** This modulation index was z-scored (see methods), and only 5.2% of cells had significant firing rate modulation beyond z of ±2.

**Extended Data Fig. 12.**
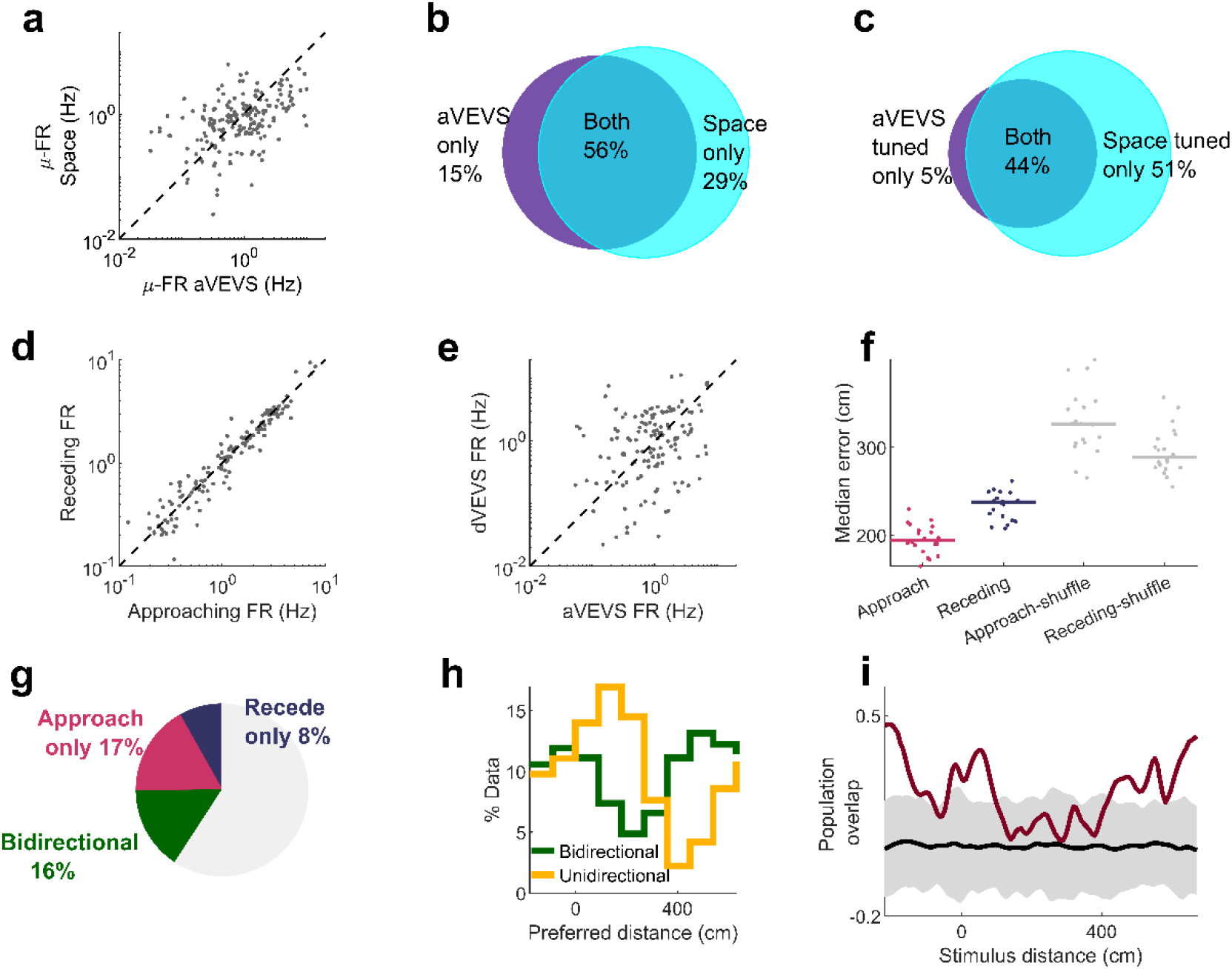
Relationship between place cells, stimulus angle (aVEVS) and distance (dVEVS) tuned cells. **(a)** The mean firing rates of cells was significantly correlated (*Pearson’s r* =0.43 *p*=4.5×10^−10^) between the aVEVS and place cell (spatial exploration) experiments. **(b)** Majority of cells active during the aVEVS experiments were also active during random foraging in real world. **(c)** Almost all of the aVEVS cells were also spatially selective during spatial exploration. **(d)** Between the approaching and receding directions, the mean firing rates, computed when the rats were immobile, were highly correlated (*Pearson’s r*=0.96 *p*=4×10^−81^) and not significantly different (KS-test *p*=0.99). **(e)** Firing rates, computed when rats were stationary, during the stimulus angle and stimulus distance experiments were significantly correlated (*r*=0.22 *p*=0.008). **(f)** Population vector decoding of the stimulus distance (similar to stimulus angle decoding, Fig 2), was significantly better than chance. (KS-test *p*=5.5×10^−10^ for approaching and *p=*4.7×10^−9^ for receding data). Approaching stimulus decoding error (median=194cm) was significantly lesser than that for receding (median=237cm) (KS-test *p*=4.2×10^−5^). These errors were 59% and 82% of the error expected from shuffled data, which was greater than that for aVEVS decoding, where the error was 33% of the shuffles, when controlling for the number of cells. **(g)** More than twice as many cells were unidirectional tuned for approaching (coming closer) movement direction, as compared to receding (moving away). **(h)** For bidirectional cells, location of peak firing in the approaching and receding direction shows bimodal response, with most cells preferring either the locations close to the rat, i.e., 0 cm or far away, ∼500cm. Unidirectional cells preferred locations close to the rat. Population vector overlap, (Fig. 5h), was further quantified by comparing the values along the diagonal for actual tuning curves, with the spike train shuffles. The actual overlap was significantly above two standard deviations of the shuffles for distances close to the rat (around 0) and far away (beyond 400cm).

**Extended Data Fig. 13.**
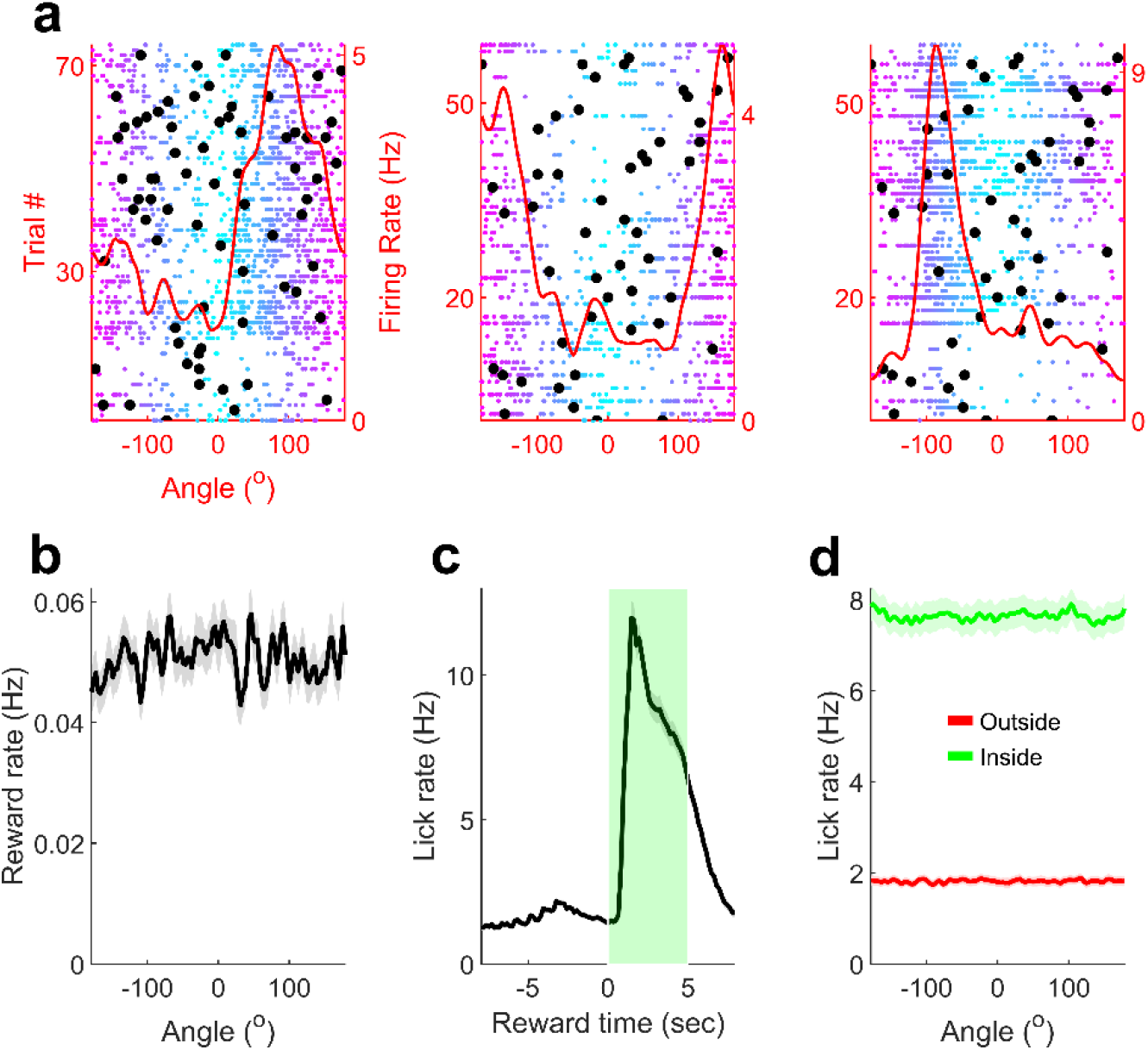
Rewards and reward related licking are uncorrelated with VEVS. **(a)** Example cells showing aVEVS from Figure 1, with reward times overlaid (black dots), showing random reward dispensing at all stimulus angles. **(b)** The average rate of rewards was uncorrelated with visual stimulus angle (circular test for uniformity *p*=0.99) **(c)** Rat’s consumption of rewards, estimated by the reward tube lick rate, was measured by an infrared detector attached to the reward tube^19^. As expected, lick rate increased after reward delivery by ∼4 fold and remained high for about five seconds (green shaded area). This duration is termed the “reward zone”. **(d)** Lick rate inside the reward zone (green) was significantly larger than that outside (red, KS-test *p*= 2.3 × 10^−54^). Inside as well as outside reward-zone lick rates were uncorrelated with visual stimulus angle (circular test for uniformity *p=*0.99 for both).

**Extended Data Fig. 14.**
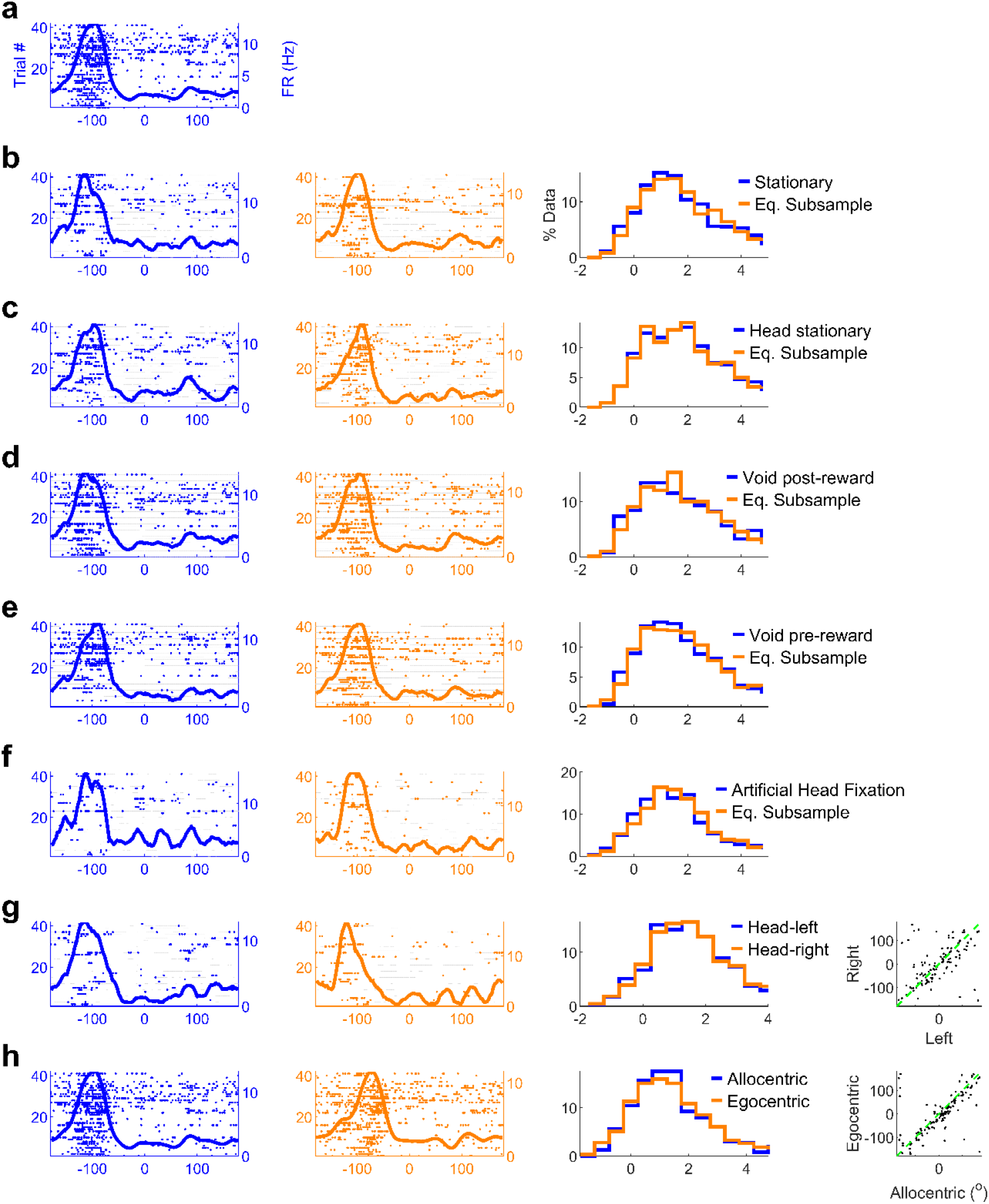
Behavioral controls of VEVS: To ascertain whether systematic changes in behavior caused VEVS, we employed a ‘behavioral clamp’ approach and estimated tuning strength using only the subset of data where the hypothesized behavioral variable was held constant. **(a)** Example aVEVS tuned cells maintained its tuning even if we used only the data when the rat was **(b)** stationary (running speed <5cm/sec, blue, left). This was comparable to a random subsample of behavior, obtained by shuffling the indices of spikes and behavior when the animal was stationary (orange, middle) (see methods). 38% of cells were aVEVS tuned (sparsity *z*>2) when using only the stationary data which is significantly greater than chance, whereas 42% were significantly tuned in the equivalent, random subsample and this difference was significant (KS-test *p=*0.02). **(c)** Similar to (b) but using only the data when the rat’s head was immobile (head movement velocity <10mm/sec). 43% and 42% of cells were significant tuned in actual behavioral clamp and equivalent subsample, and these were not significantly different (KS-test *p=*0.93) **(d)** Similar to (b), but removing data within 5 seconds after reward dispensing, called void post-reward. 43% cells were tuned in “void post-reward” data, 43% for equivalent subsample (KS-test *p*=0.56). **(e)** Similar to (d), but removing data within 5 seconds before reward dispensing, called void pre-reward. 39% cells were tuned for void pre-reward, 42% for equivalent subsample (KS-test *p*=0.43). **(f)** Using a subsample of data, from when the rat’s head was within the central 20 percentile of head positions (typically <10°), rat was stationary and there were no rewards in the last 5 seconds. This condition was called “analytical head fixation”. 28% of cells were aVEVS tuned under this behavioral clamp, which was lesser than that in an equivalent subsample (31%, KS-test *p*=0.05), but significantly greater than chance. **(g)** Tuning curves for head positions to the leftmost 20 percentile and rightmost 20 percentile were similar, with 31% and 32% cells tuned in the two conditions (KS-test *p*=0.67). The preferred angles of tuning were highly correlated (circular correlation *r*=0.67 p=1.3×10^−11^) and not significantly different (circular KS-test *p>0*.*1*). **(h)** aVEVS tuning was recomputed in the head centric frame, by accounting for the rat’s head movements (obtained by tracking overhead LEDs attached to the cranial implant) and obtaining a relative stimulus angle, with respect to the body centric head angle. Overall tuning levels were comparable, between allocentric and this head centric estimation. First panel of (h) is the same as that in (a) since all aVEVS tuning reported earlier was in the allocentric or body centric frame. Using a subset of data when both overhead LEDs were reliably detected, 25% and 26% of cells were significantly tuned for the stimulus angle in the allocentric and egocentric frames (KS-test *p*=0.9). Preferred angle of aVEVS tuning for tuned cells was highly correlated (*r*=0.81, *p* = 1.8×10^−15^) and not significantly different between the two frames (circular KS-test *p*>0.1).

**Extended Data Fig. 15.**
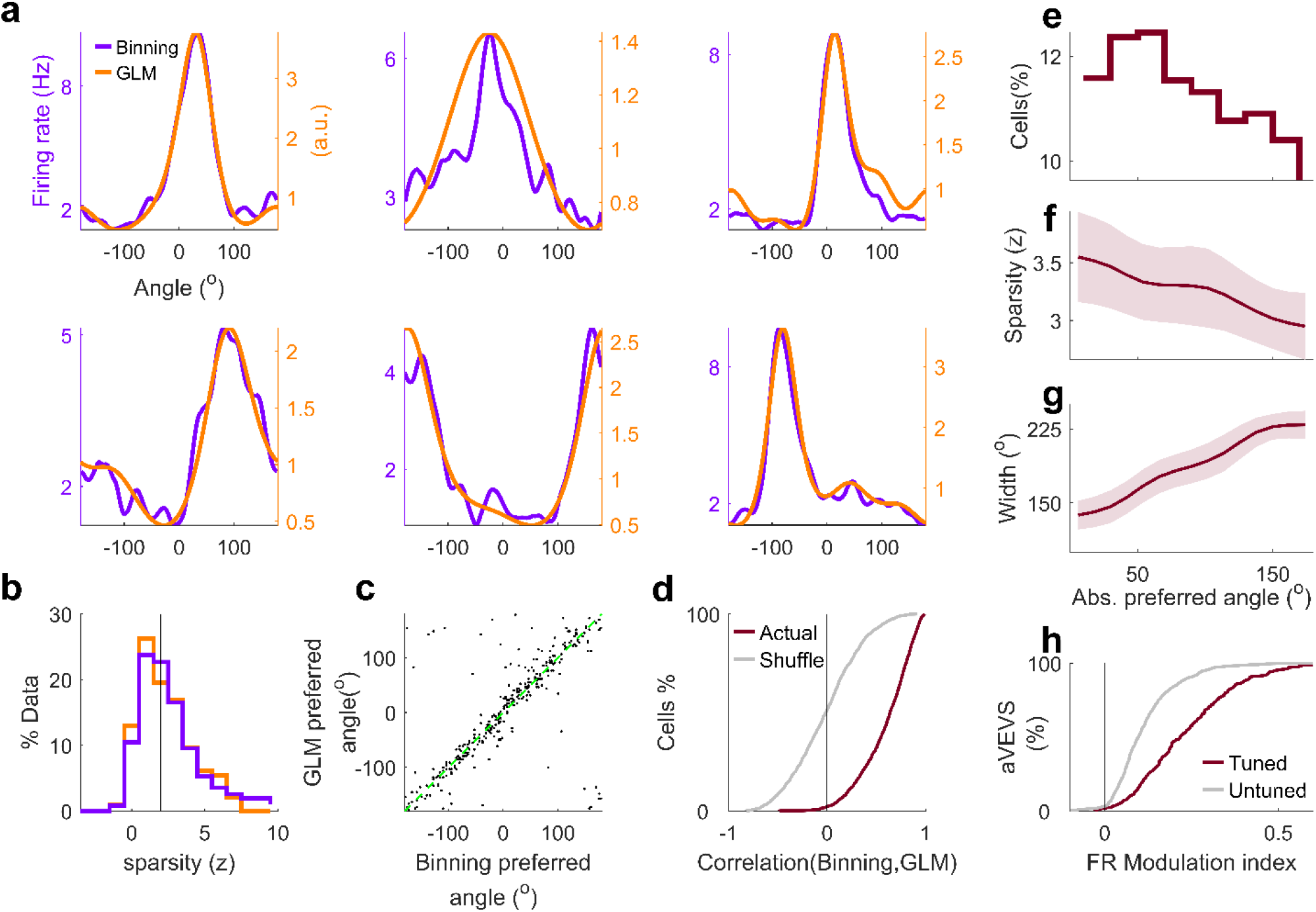
GLM estimate of aVEVS tuning. To estimates the independent contribution of stimulus angle to neural activity, while factoring out the contribution of head position and running speed, we used the generalized linear model (GLM) technique (see methods)^18^. **(a)** Tuning curves obtained by binning methods were comparable with those from GLM estimation, including for the cells used in Fig. 1 (first 2 examples in row 1 & 2). **(b)** Sparsity levels were comparable (KS-test *p*=0.07) and 40% of cells were found to be significantly tuned for stimulus angle using GLM based estimated, compared to 43% from binning in this subset of data where head and leg movements were reliably captured (cell count, n=991). **(c)** Preferred angle of firing between GLM and binning based estimates of aVEVS were highly correlated (circular correlation test *r=*0.86 *p*<10^−150^) **(d)** Correlation between the aVEVS tuning curves from the two methods was significantly greater than that expected by chance, computed by randomly shuffling the pairing of cell ID across binning and GLM (KS-test *p*<10^−150^). **(e-h)** Properties of aVEVS tuning responses based on GLM estimates were similar to those based on binning method, as shown in Figure 1. (**e**) Distribution of tuned cells as a function of the preferred angle (angle of maximal firing). There were more tuned cells at forward angles than behind. (**f**) Median ± SEM z-scored sparsity and its variability (SEM, shaded area, here and subsequently) of tuned cells as a function of their preferred angle. (*Pearson’s r*=-0.17 *p*=0.004). (**g**) Median ± SEM full width at quarter maxima across the ensemble of tuned responses increased as a function of preferred angle of tuning. (*Pearson’s r*=+0.33 *p*<10^−150^). (**h**) CDF of firing rate modulation index within versus outside the preferred zone (see methods) for tuned cells were significantly different (Two-sample KS test *p*=2.9×10^−37^).

**Extended Data Fig. 16.**
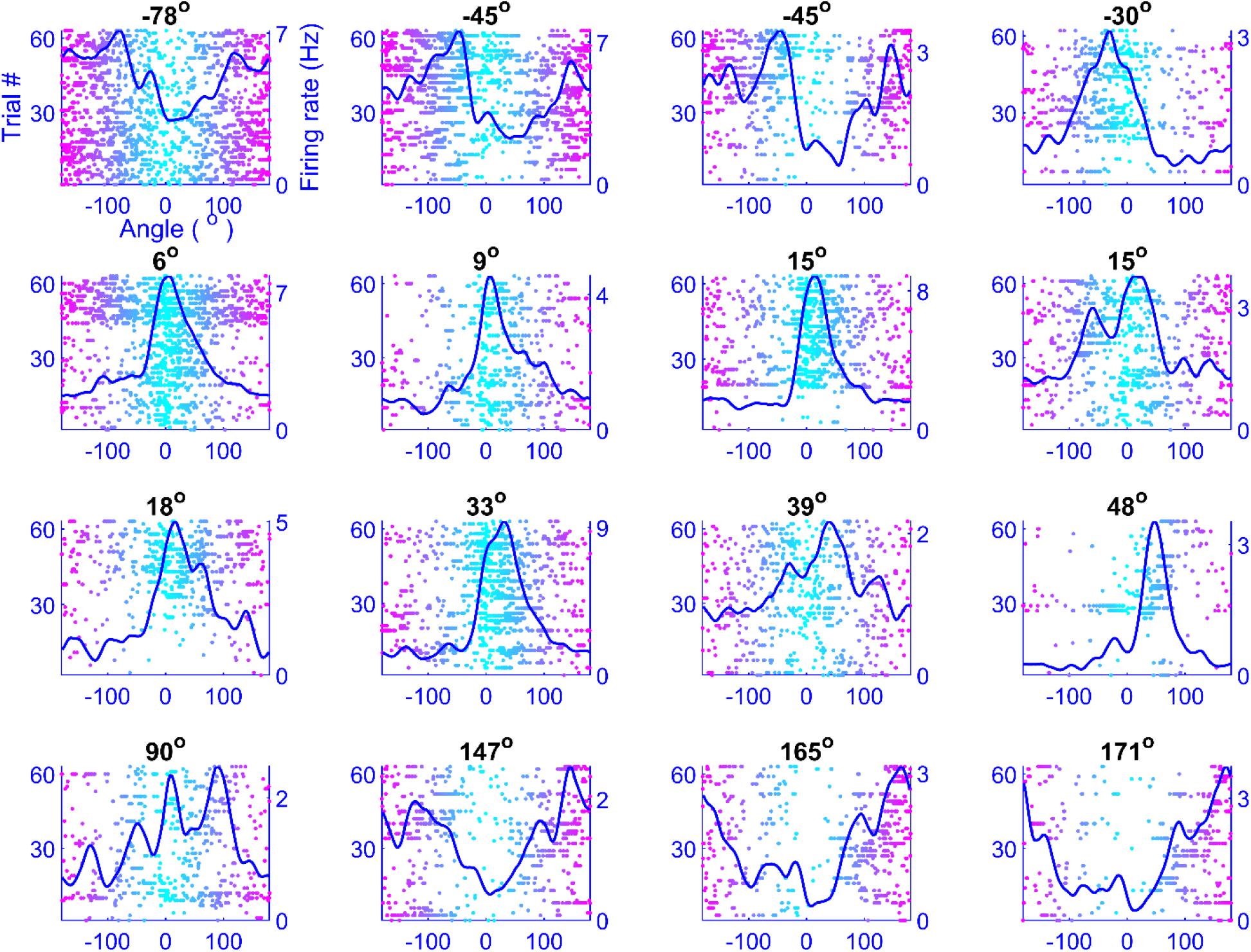
Simultaneously recoded cells span all angles, including behind the rat. 16 simultaneously recorded cells showed significant aVEVS. Their preferred angles are indicated on top. Only cells selective for CCW direction shown for clarity. While the forward direction (0°) is overrepresented, these cells span all angles of the visual field including angles behind him (180°).

**Extended Data Fig. 17.**
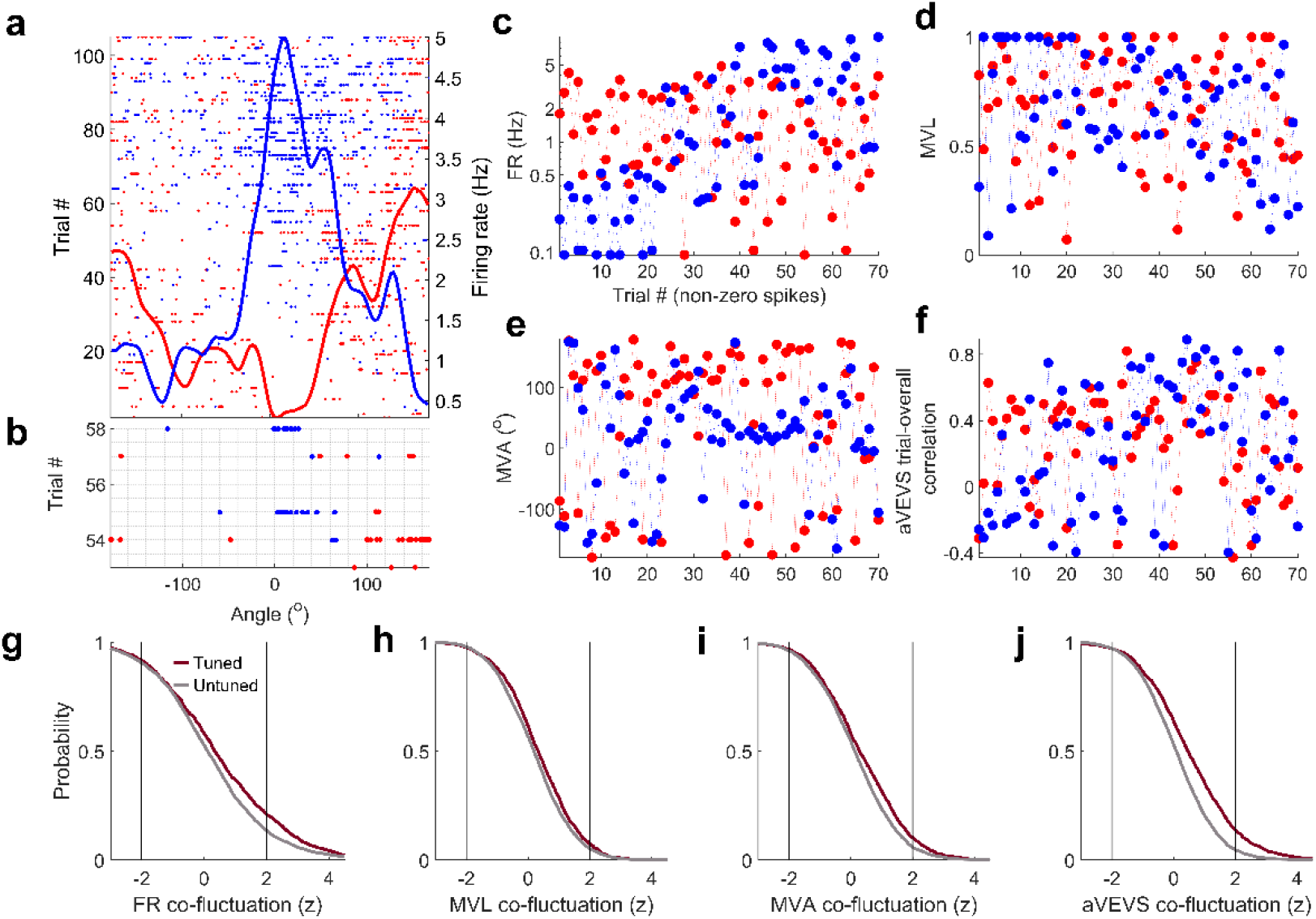
Simultaneously recorded cells show very weak co-fluctuation of aVEVS tuning across trials. **(a)** Two simultaneously recorded cells showing significant aVEVS in the CCW direction, **(b)** zoomed in for a subset of trials, showing mostly uncorrelated fluctuations in the two cells’ spiking. **(c)** For the same cell-pair, mean firing rate across trials was broadly uncorrelated. Only trials with non-zero spikes were used here, and henceforth, to ensure comparison with aVEVS tuning (see below). **(d)** Same as (c) but showing uncorrelated fluctuations in the depth of modulation of aVEVS response of the two cells across trials, quantified by the Mean Vector Length (MVL, see *Methods*). **(e)** Same as (c) but showing uncorrelated fluctuation of aVEVS response across trials, quantified by Mean Vector Angles (MVA, see *Methods*). **(f)** Same as (c) but showing largely independent fluctuations in the overall aVEVS tuning (measured by correlation between the trial-averaged aVEVS tuning curve and the aVEVS tuning curve in a given trial) for this cell-pair. The significance of co-fluctuations in cell-pairs were quantified by bootstrapping methods, by employing trial id shuffles (see *Methods*). CCW and CW tuning curves were treated as separate responses throughout these analyses. **(g)** 21% (14%) of simultaneously recorded, tuned (untuned) cell-pairs showed significant (z>2) co-fluctuation of mean firing rates across trials which provides an estimate of the non-specific effects such as running, reward consumption etc. **(h)** Only 7% (5%) of tuned (untuned) cell pairs showed significant co-fluctuation of MVL across trials indicating little effect of nonspecific variables on the depth of aVEVS tuning. **(i)** Similarly, only 10% (6%) of tuned (untuned) cell pairs showed significant co-fluctuation of MVA across trials. **(j)** Only 14% (5%) of tuned (untuned) cell pairs showed significant co-fluctuation of aVEVS. Notably, the number of cell pairs showing significant co-fluctuations in any of the aVEVS tuning properties (h, i, j) was smaller than the number of cell pairs showing significant co-fluctuation of firing rates; and there was little qualitative difference between the significantly aVEVS tuned vs untuned populations.

